# Estimating maximal microbial growth rates from cultures, metagenomes, and single cells via codon usage patterns

**DOI:** 10.1101/2020.07.25.221176

**Authors:** Jake L. Weissman, Shengwei Hou, Jed A. Fuhrman

## Abstract

Maximal growth rate is a basic parameter of microbial lifestyle that varies over several orders of magnitude, with doubling times ranging from a matter of minutes to multiple days. Growth rates are typically measured using laboratory culture experiments. Yet, we lack sufficient understanding of the physiology of most microbes to design appropriate culture conditions for them, severely limiting our ability to assess the global diversity of microbial growth rates. Genomic estimators of maximal growth rate provide a practical solution to survey the distribution of microbial growth potential, regardless of cultivation status. We developed an improved maximal growth rate estimator, and implement this estimator in an easy-to-use R package (gRodon), which outperforms the state-of-the-art growth estimator in multiple settings, including in a community context where we implement a novel species abundance correction for metagenomes. Additionally, we estimate maximal growth rates from over 200,000 genomes, metagenome-assembled genomes, and single-cell amplified genomes to survey growth potential across the range of prokaryotic diversity. We provide these compiled maximal growth rates in a publicly-available database (EGGO), which we use to illustrate how culture collections show a strong bias towards organisms capable of rapid growth. We demonstrate how this database can be used to propagate maximal growth rate predictions to organisms for which we lack genomic information, on the basis of 16S rRNA sequence alone. Finally, we observe a bias in growth predictions for extremely slow-growing organisms, ultimately leading us to suggest a novel evolutionary definition of oligotrophy based on the selective regime an organism occupies.

**Significance:** Despite the wide perception that microbes have rapid growth rates, many environments like seawater and soil are often dominated by microorganisms that can only grow very slowly. Our knowledge about growth is necessarily biased towards easily culturable organisms, which turn out to be those that tend to grow fast, because microbial growth rates have traditionally been measured using lab growth experiments. But how are potential growth rates distributed in nature? We developed a tool to predict maximum growth rate from an organism’s genome sequence (gRodon). We predicted the growth rates of over 200,000 organisms and compiled these predictions in a publicly-available database (EGGO), which illustrates how current collections of cultured microbes are strongly biased towards fast-growing organisms.

## Introduction

Microbial growth rates vary widely, with doubling times ranging from under 10 minutes for lab-reared organisms [15] to several days for oligotrophic marine organisms [39, 52], and even as high as many years for deep sub-surface microbes [11, 60, 67]. Even under optimal nutrient conditions and in the absence of competition, species will vary in their maximal potential growth rates as a function of their ability to rapidly synthesize cellular components and replicate their genome [33, 28, 72, 55]. Broad lifestyle differences can be detected across habitats, with many oligotrophic marine systems harboring slow-growing organisms relative to nutrient-rich habitats like the human gut [72, 62]. Yet, optimal, or even adequate, culture conditions for the majority of prokaryotic organisms are unknown [53, 25], making it difficult to assess the true diversity of microbial maximal growth rates. Although growth media for some species can be predicted based on their phylogeny [44], cultivation is laborious and impractical in a high-throughput manner for many ecosystems such as deep sea waters. Moreover, as we show here, even comprehensive culturing efforts targeted at a specific ecosystem (e.g., the human gut) tend to be biased towards fast-growing members of the community. By estimating maximal growth rates directly from environmentally-derived sequences it may be possible to build a comprehensive and unbiased snapshot of growth across different habitats.

A beacon of hope, maximal growth rates predicted using genome-wide codon usage statistics [72] appear to capture overall trends in the growth rates of natural communities [36]. Because the genetic code is degenerate, genes may vary in their usage of alternative codons for a given amino acid. Highly expressed genes demonstrate a biased usage of alternative codons, optimized to cellular t-RNA pools [26, 21, 14, 24, 63, 18]. Vieira-Silva et al. [72] showed that among several possible genomic indicators of growth (e.g., rRNA copy number and proximity to the origin of replication, t-RNA copy number, etc.) high codon usage bias (CUB) in genes coding for ribosomal proteins and other highly-expressed genes is the best predictor of high maximal growth rates, and can be used to make accurate predictions even with partial genomic data. Their growthpred software leverages this bias to predict maximal growth rates from genomic data [72].

We extend the work of Vieira-Silva et al. [72] by assessing additional dimensions of codon usage [63, 10]. In doing so we are able to substantially improve our predictive performance. Additionally, we provide a correction based on species abundances to the method when applied to bulk community data from metagenomes, an important but previously neglected correction. Together we provide a user-friendly implementation of these methods in an R package (gRodon). Using our method, we assay growth rates in over 200,000 genomes ([65, 66, 23]) and environmentally-derived metagenome-assembled genomes (MAGs; [48, 69, 61, 1, 74]) and single-cell amplified genomes (SAGs; [8, 46]) in order to survey the natural diversity of prokaryotic growth rates. We provide this comprehensive set of over 200,000 predictions as a compiled database of estimated growth rates (estimated <u>g</u>rowth rates from <u>g</u>Rodon <u>o</u>nline; EGGO). Using this large database we demonstrate how growth rate predictions can be propagated to organisms for which no genomic information is available but that have a close relative in EGGO. Finally, we provide guidance as to when codon-usage based growth estimators are expected to fail, and when classification (i.e. predicting oligotrophy vs. copiotrophy) may be a wiser use of these methods than regression (i.e., prediction of exact doubling times).

## Results and Discussion

### Predicting Maximal Growth Rates

#### More than one aspect of codon usage is associated with growth

We measured three features of codon usage: (1) the median CUB of a user-provided set of highly-expressed genes relative to the codon usage pattern of all genes in a genome [63], (2) the mean of the CUBs of each highly-expressed gene relative to the overall codon usage pattern of the entire set of highly-expressed genes, and (3) the genome-wide codon pair bias [10]. For details of these calculations see the Methods. In practice, we take the set of highly-expressed genes to be those coding for ribosomal proteins because these genes are expected to be highly expressed in most organisms [72]. The first (1) measure captures CUB in the classical sense, and the MILC metric we use [63] controls for overall genome composition as well as gene length. The second (2) measure captures the “consistency” of bias across highly expressed genes, with the intuition that if highly-expressed genes are optimized to cellular t-RNA pools then they will share a common bias (low values indicate high consistency). This quantity can be though of as the “distance” between highly expressed genes in codon-usage space, where we expect these genes to be close together when they are highly optimized for growth. The third (3) measure, codon pair bias, captures associations between neighboring codons, which have been suggested to impact translation [22, 6, 10]. Specifically, it has been shown that altering the frequency of different codon pairs (but not the overall codon or amino acid usage) can lead to lower rates of translation, and this strategy has been used to produce attenuated polioviruses (potentially to engineer novel vaccines; [10]). Because it is much more difficult to accurately estimate pair-bias due to the large number of possible codon pairs, we do so on a genome-wide scale, calculating pair-bias over all genes rather than just for highly expressed genes (our R package includes a “partial” mode for when this is not possible due to partial genomic information). Consider that if there are 64 codons, the number of possible ordered pairs is 4096, and accordingly far more data will be needed to reliably estimate the frequencies of all of these pairs than the original set of codons.

We fit our model using all available completely assembled genomes in RefSeq (1415) for the set of 214 species with documented maximal growth rates compiled by Vieira-Silva et al [72]. All three of these measures were significantly associated with growth rate in a multiple regression (CUB, *p* = 2.2 *×* 10^*-*37^; consistency, *p* = 8.1 *×* 10^*-*15^; codon-pair bias, *p* = 5.3 *×* 10^*-*6^; linear regression). Furthermore, comparing nested models, incorporating first CUB, then consistency, and finally codon-pair bias, we found that each nested model fit the data significantly better than the last (addition of consistency, *p* = 4.2 *×* 10^*-*11^; addition of codon-pair bias, *p* = 4.0 *×* 10^*-*6^; likelihood-ratio test).

### gRodon accurately predicts maximal growth rates

The gRodon model fit the available maximal growth rate data well (adjusted *R*^2^ = 0.63; Fig 1a). Our model demonstrated a significantly better fit to growth data than a linear model fit on the output of growthpred (ANOVA, *p* = 1.1 *×* 10^*-*8^; Fig 2). Notably, gRodon provided a better fit to the data than growthpred at both high and low growth rates (S1 Figure).

**Figure 1:**
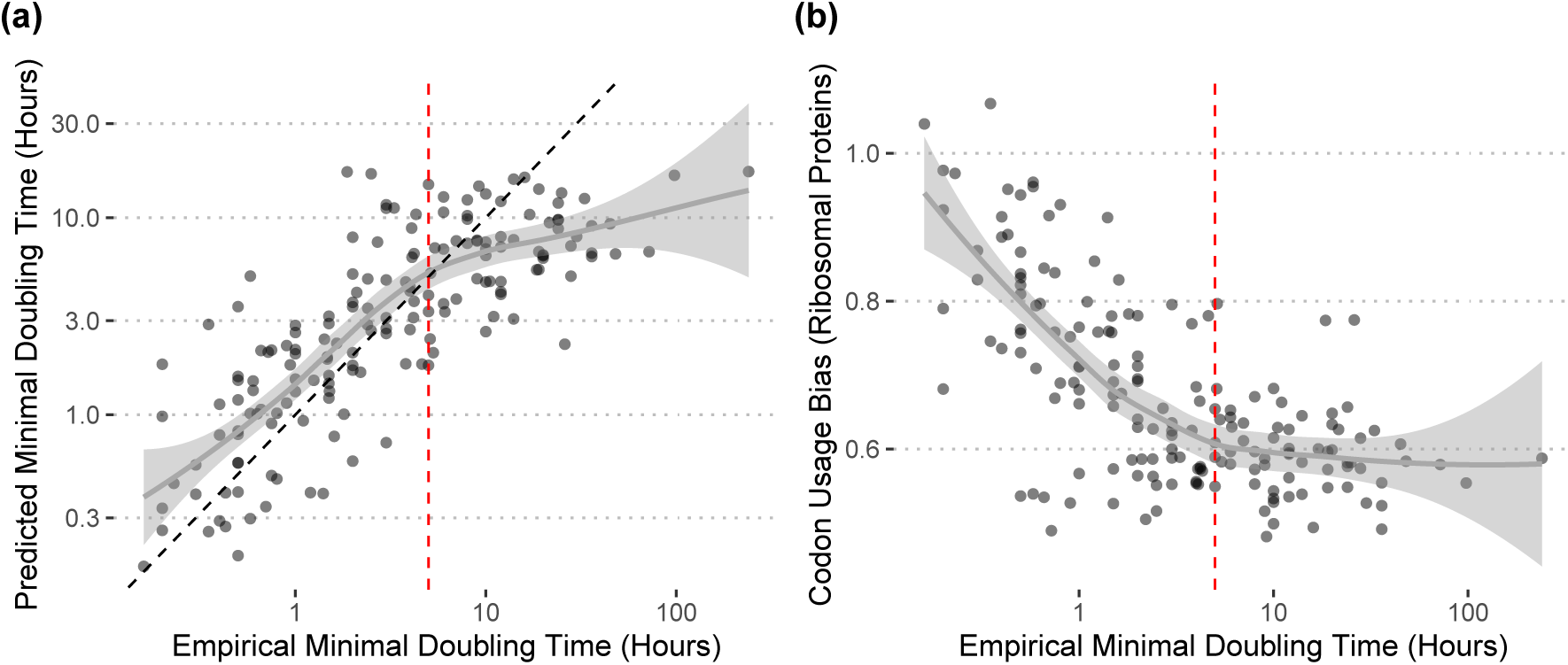
Predictions from gRodon accurately reflect prokaryotic growth rates, with the caveat that (a) gRodon underestimates doubling times when growth is very slow due to (b) a floor on CUB reached in slow-growth regimes. Vertical dashed red line at 5 hours indicates where the CUB vs. doubling time relationship appears to flatten. The black dashed line in (a) is the *x* = *y* reference line.

**Figure 2:**
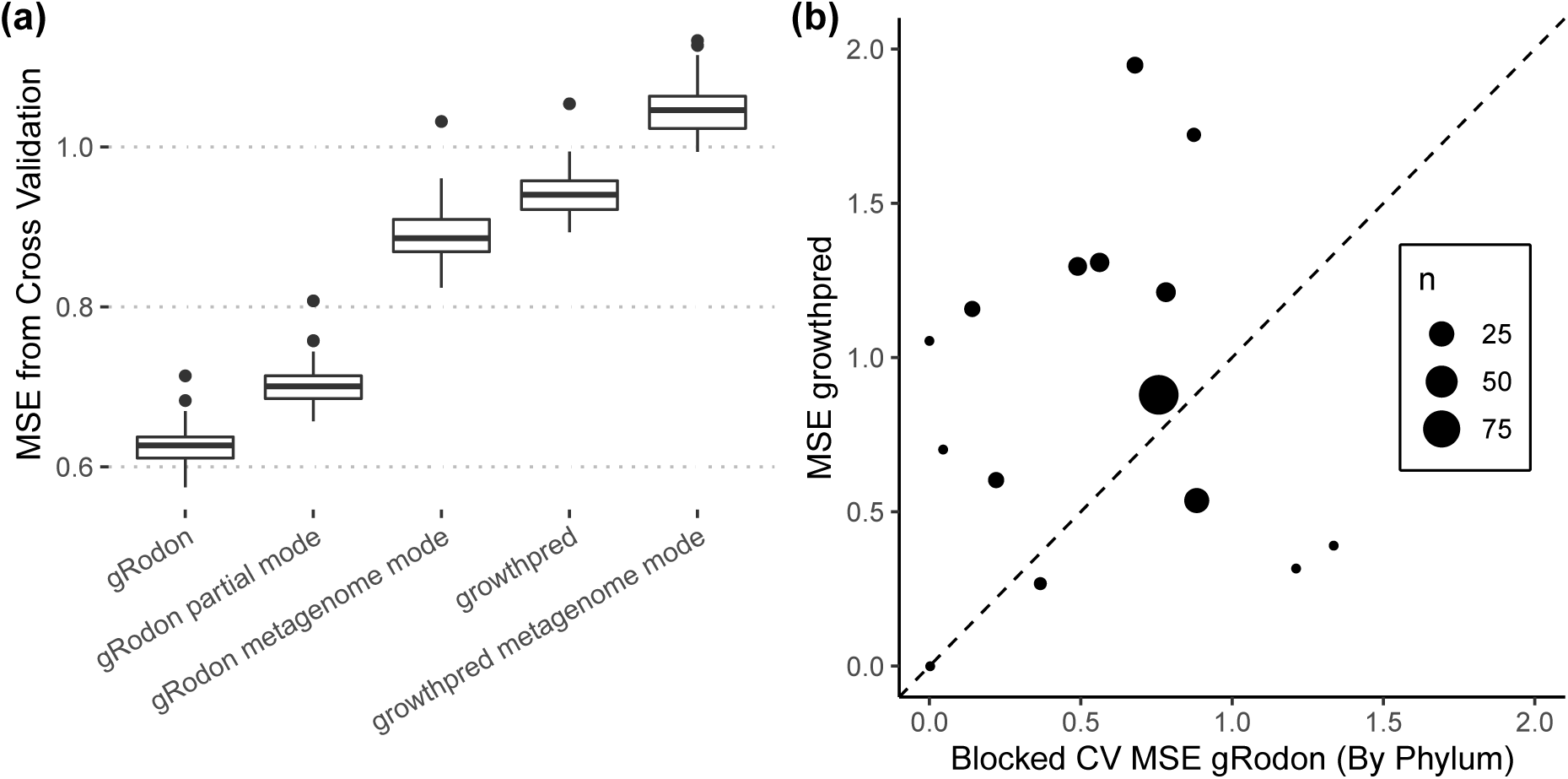
Predictions from gRodon are more accurate that those from growthpred. (a) Under 10-fold cross-validation (CV; repeated 100 times) gRodon outperforms growthpred (in terms of mean squared error, MSE). (b) Even extrapolating across phyla gRodon typically outperforms growthpred. Each point represents error extrapolating to a given phylum, with the point size representing the number of species assigned to that phylum in our dataset. The black dashed line is the *x* = *y* reference line. Note that in both (a,b) the growthpred values shown are not cross-validated (since growthpred’s model has already been fit on the full dataset), but performance values were calculated on each fold, giving growthpred an advantage (though gRodon still demonstrates higher accuracy despite the unfair comparison).

We considered the possibility of overfitting our model to the data, which would inhibit our ability to apply our predictor to new datasets. Overfitting is a particularly relevant concern when dealing with species data, since models may end up being fit to underlying phylogenetic structure rather than real associations between variables. In addition to traditional cross-validation (Fig 2a), we implemented a blocked cross validation approach, which effectively controls for phylogenetic structure when estimating model error [54]. Under this framework, we take each phylum in our dataset as a fold to hold out for independent error estimation rather than holding out random subsets of our data as in traditional cross validation. We found that even when predicting growth rates for each phylum in this way (extrapolating from our model fit to all other phyla, but excluding the test phylum), we outperformed growthpred’s predictions for the large majority of phyla (Fig 2b). Importantly, for this comparison growthpred’s predictions were based on it’s fit to the entire dataset (including the test phylum), meaning that gRodon was able to outperform growthpred even when given an unfair disadvantage.

We examined a number of confounding variables that could affect model performance. Observed codon statistics are the result of several interacting evolutionary forces. Selection for rapid growth drives the signal we exploit here, but the effective population size (*N*_*e*_) and the rate of recombination will determine how efficiently selection acts on a given population [12]. We found that *N*_*e*_ is correlated with maximal growth rate (as might be expected; [3]), as well as our model residuals (S2 Figure), though the effect is rather weak. For populations with extremely atypical effective population sizes (e.g., intracellular symbionts), we caution that *N*_*e*_ is likely to confound genomic growth rate estimates. Recombination locally increases the efficiency of selection, and can lead to weak but significant patterns in GC content along the genome [2, 73]. We found no apparent differences in codon usage bias between genes with or without a signal of recombination, both looking at all genes in a genome (S3 Figure) and just the ribosomal proteins (S4 Figure). Finally, especially in oligotrophic marine environments, many microbes experience selection for genome streamlining (high percent coding sequence) alongside selection for low genomic GC content [64, 20]. While our measures of codon usage should correct for genome nucleotide composition, we wanted to be sure our model’s performance was not affected by these other targets of selection. While percent coding sequence does appear to have some non-linear association with growth rate, our model residuals were not affected by either percent coding sequence or GC content (S5 Figure). This is consistent with previous work showing that CUB-based approaches can predict growth rates in low-nutrient marine microcosms [36].

Finally, we assessed the impact of our training set on gRodon’s predictions. The original set of minimal doubling times from Vieira-Silva et al. [72] was a carefully hand-curated dataset compiled specifically for this application, but includes only a subset of available recorded doubling time estimates for cultured microbes. Unfortunately, there is no single database describing all known microbial growth rates, but recent work has attempted to compile all available microbial phenotypic data [38], including data on growth rates. We re-trained gRodon on the growth rates associated with microbes with completely assembled genomes in the Madin et al. [38] database (464 species with 8287 genomes). The re-trained model yields very similar results to the original gRodon model (S6 and S7 Figures), despite the two training datasets disagreeing on the maximal growth rates of several species (S6 Figure). We include this alternative model in the gRodon package alongside the model trained only on the Vieira-Silva et al. [72] dataset and include predictions from both models for each entry in the EGGO database.

### The problem of slow-growers

For very long doubling times, while gRodon outperforms growthpred it still tends to underestimate the actual doubling time (Fig 1a and Fig 2a). In populations of very slow growing microbes, selection to optimize transcription of ribosomal proteins is likely quite low, and once the selective coefficient is low enough, drift will dominate the evolutionary process. This expectation is consistent with the pattern seen in Fig 1b where CUB of the ribosomal proteins reaches a floor at very high doubling times. Importantly, this floor will likely be a problem for all genomic predictors of maximal growth rate. Drift will be the primary evolutionary force influencing any genomic feature when selection coefficients approach zero, as we expect for genomic features associated with rapid growth in extremely slow-growing organisms.

What can be done in such a scenario? While gRodon cannot accurately differentiate between a doubling time of 10 or 100 hours, it can reliably tell us if a doubling time is greater than 5 hours long (the threshold at which CUB flattens in Fig 1b, see S8 Fig). In fact, this threshold suggests a natural definition of an oligotroph as an organism for which selection for rapid maximal growth is low enough so that no signal of growth optimization (e.g., CUB) is observed. Importantly, this standard redefines oligotrophy in evolutionary terms, as a specific selective regime that a microbe can occupy, and therefore the threshold for oligotrophy will depend on the *N*_*e*_ of a species (as illustrated by the effects of *N*_*e*_ on our model residuals above). From our data, it appears that at typical *N*_*e*_ values for microbes (*∼* 10^8^; [3], S2 Fig), codon optimization is undetectable for maximal doubling times greater than 5 hours (Fig 1b and S8 Fig). Even for Prochlorococcus marinus, which may have very large effective population sizes (*>* 10^13^ [27] over a well-mixed marine region, though some estimates of Prochlorococcus *N*_*e*_ are much lower at *∼*10^8^ [3]), growth rates were severely underestimated, though still above our 5 hour threshold (predicted doubling time of 6.2 hours versus an actual doubling time of 17 hours for strain CCMP1375). Thus, gRodon can be used as an accurate classifier for oligotrophy/copiotrophy by simply defining microbes predicted to have maximal doubling times greater than 5 hours as oligotrophs (S8 Fig). Obviously this threshold will vary to some degree across species and populations (e.g, as local population size, population structure, selective regimes, recombination rates, etc. vary), but our predictor appears to be largely robust to most confounders (S2, S3, S4, and S5 Figures), and without additional information 5 hours serves well as a pragmatic default.

### Predicting the mean growth rate of a community using metagenomes

We cannot resolve the genomes of the majority of organisms described by a typical metagenomic sample. Yet, often we wish to look for changes in community-scale characteristics over space and time. Given a nearly complete set of coding sequences from a community, is it possible to estimate community-wide growth potential even when we do not know which organisms make up that community? Vieira-Silva et al. [72] found differences in the CUB across habitats and during ecological succession in the infant gut, interpreting this as community-level differences in the average maximal growth rate. This approach is supported by the fact that codon usage patterns and t-RNA copy numbers tend to be shared by members of a community [72, 68, 56], where different species within an environment tend to have more similar codon usage patterns than the same species in different environments [56]. Thus, comparing the set of all highly expressed genes (e.g., all genes coding for ribosomal proteins) to the full set of genes in a metagenome should give a rough estimate of the mean community-wide growth rate.

Importantly, the growthpred approach makes a major omission in that it does not account for the relative abundances of different organisms in the sample. All assembled genes are treated as equal, thus biasing the growth estimate towards the rarer members of a community. To correct for this, we incorporated read coverage of genes into our gRodon calculation, thus producing a community-wide maximal growth rate estimate that reflects the taxonomic distribution of a community. Our approach is simple - in gRodon’s metagenome mode (which only takes CUB into account, not consistency or pair-bias) we calculate the weighted median of the CUB of highly expressed genes, with weights corresponding to the mean depth of coverage of these genes, rather than an unweighted median as in the default gRodon calculation. Thus, the highly expressed genes of abundant organisms are accounted for proportionally to their relative abundance. For comparison, we also implemented an unweighted version of metagenome mode in gRodon.

In practice, it is not easy to benchmark such a method on a natural community since we do not typically know the actual maximal growth rates of all members of any given community. Nevertheless, our approach can be validated by nutrient enrichment experiments where nutrients are added to an initially oligotrophic community leading to a rise in copiotrophs. If gRodon truly captures changes in community-wide growth potential, we should see our community-level maximal growth rate predictions increase under this nutrient enrichment regime. While many such experiments have been carried out, very few are accompanied by shotgun metagenomic sequencing. Recently, Okie et al. [45] performed a controlled nutrient enrichment experiment in a highly oligotrophic pond system that included replicated metagenomic samples from the treatment and control conditions. Despite a small number of samples overall (*n* = 10), gRodon’s weighted metagenome mode detected a significantly higher community-level average maximal growth rate in the enrichment condition (*p* = 0.032, Mann-Whitney test; S10 Fig). Importantly, no difference was detected when using gRodon’s unweighted metagenome mode (*p* = 0.15, Mann-Whitney test; S10 Fig). Okie et al. [45] excluded several samples from their final analyses on the basis of low read counts, doing the same sufficiently reduces our sample size (*n* = 7) so that no significant change is detected (*p* = 0.057, Mann-Whitney test), though all enriched treatments have higher predicted maximal growth rates than all control treatments (S11 Fig). In a recent time-series study, Coella-Camba et al. [9] applied multiple nutrient treatments to mesocosms in oligotrophic marine waters and tracked their change over time with shotgun metagenomes. In several experiments a large cyanobacterial bloom was observed within the first 7 days of the experiment followed by a crash [9], which both gRodon’s weighted and unweighted metagenome modes were able to capture as a steep increase in growth rate before a return to baseline (S12 Fig), though the un-corrected, unweighted metagenome mode systematically underestimated average community maximal growth rates (S13 Fig). As sequencing costs continue to decline it will become easier to benchmark community-wide maximal growth estimates, though even from our limited example we emphasize that it is important to take relative abundances into account when making these estimates.

### The EGGO Database

We constructed a database (EGGO; Table 1) of predicted growth rates from 217,074 publicly available genomes, metagenome-assembled genomes (MAGs), and single-cell amplified genomes (SAGs). Of these, the majority corresponded to RefSeq genome assemblies (184,907; [65, 66]). The distribution of growth rates across RefSeq was roughly bi-modal, with the split between peaks corresponding to the 5 hour doubling-time cutoff we proposed above for classifying oligotrophs (Fig 3a). Additionally, phyla tended to broadly group together in terms of growth rate, and the 5 hour divide separated fast- and slow-growing phyla (Fig 3b-c). Using a Gaussian mixture model we obtained two large clusters of microbes, with mean doubling times of 2.7 and 7.9 hours respectively, roughly corresponding to our proposed copiotroph/oligotroph divide (Fig 3a). We also obtained a third, very small and slow growing cluster, accounting for 0.4% of observations with a mean minimal doubling time of 99 hours (too small to plot in Fig 3a).

**Table 1:** Summary of EGGO database

| Source | Type | Number of Genomes | Environment |
| --- | --- | --- | --- |
| RefSeq Assemblies [65] | Isolate | 184907 | - |
| Parks et al. [48] | MAG | 7287 | - |
| GORG-tropics [46] | SAG | 7214 | Marine Surface |
| Tully et al. [69] | MAG | 2266 | Marine |
| Delmont et al. [13] | MAG | 809 | Marine |
| MarRef [29] | Isolate | 725 | Marine |
| Pasolli et al. [49] | MAG | 4431 | Human Microbiome |
| Nayfach et al. [41] | MAG | 4483 | Human Gut |
| Poyet et al. [50] | Isolate | 3459 | Human Gut |
| Zou et al. [75] | Isolate | 1493 | Human Gut |

**Figure 3:**
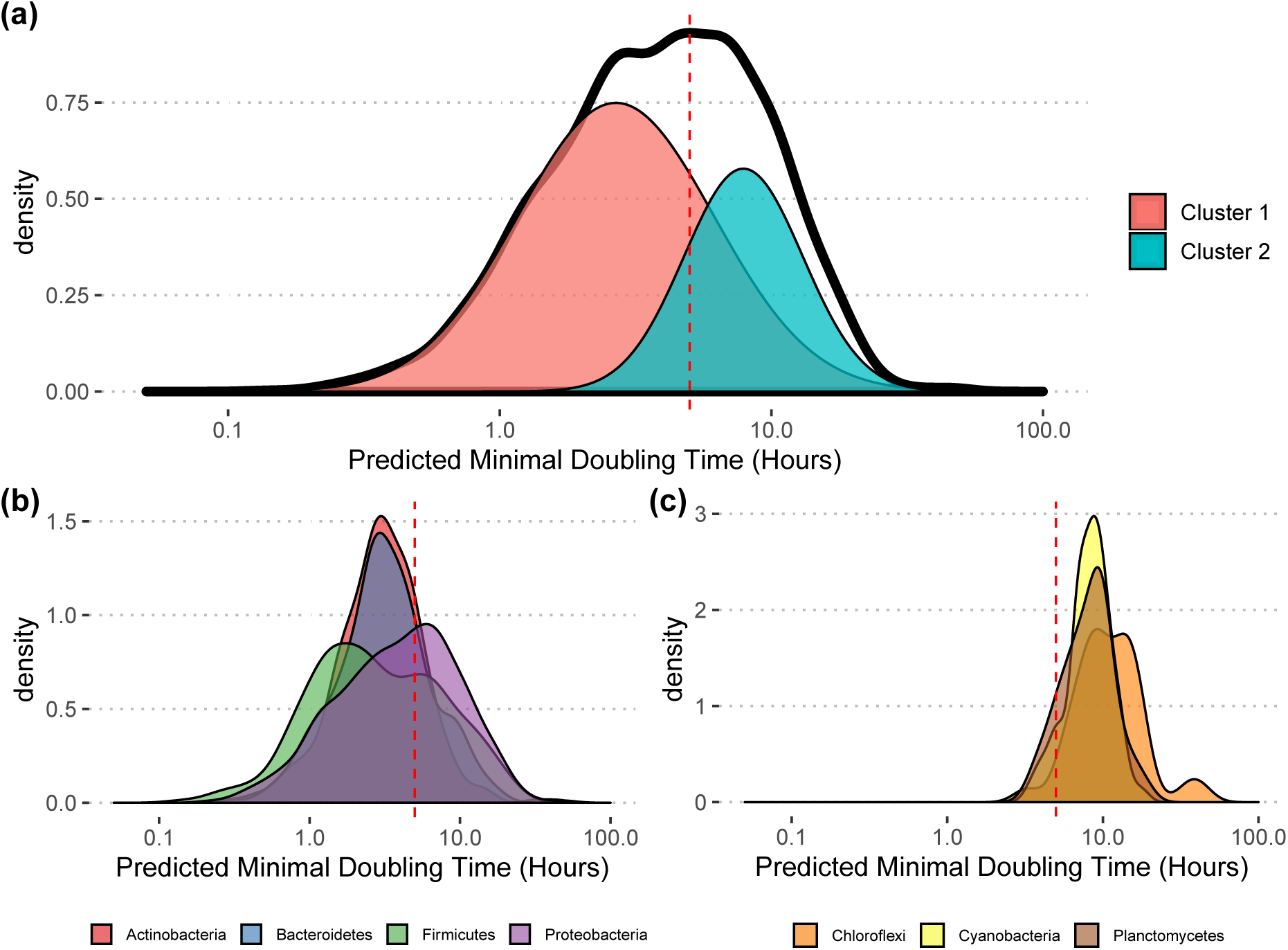
Prokaryotes with sequenced genomes span a broad range of predicted growth rates. (a) Predicted growth rates for assemblies in NCBI’s RefSeq database. Growth rates were averaged over genera to produce this distribution, since the sampling of taxa in RefSeq is highly uneven (see S9 Fig for full distribution; a small number of genera had inferred doubling times over 100 hours, 6 out of 2984). Clusters correspond the components of a Gaussian mixture model, with area under each curve scaled to the relative likelihood of an observation being drawn from that cluster. (b-c) Growth rate distributions for individual (b) fast- and (c) slow-growing phyla (only showing phyla with *≥* 30 genera represented in RefSeq). Vertical dashed red line in (a-c) at 5 hours for reference.

MAGs and SAGs make up a sizable portion of our overall database (26,490) and provide important information about the distribution of growth rates of uncultured organisms. A basic expectation is that cultured microbes from an environment will on average have higher maximal growth rates than the true average across that environment, since culturing slow-growing species will in general be more difficult [53, 70]. This pattern can be clearly seen in both marine (Fig 4) and host-associated (Fig 5a-b) environments, with isolate collections showing much lower predicted doubling times than MAGs and SAGs from the same environments. Even in sets of isolates meant to capture the complete taxonomic diversity in an environment [75, 50], we see that they fail to capture the most slowly-growing members of the community (Fig 5a-b). Illustrating this gap is important, as it shows how existing culture collections are not only incomplete, but also biased. These patterns are most apparent when looking within an environment, and largely disappear when comparing against MAGs from diverse environments (S14 Fig; [48]).

**Figure 4:**
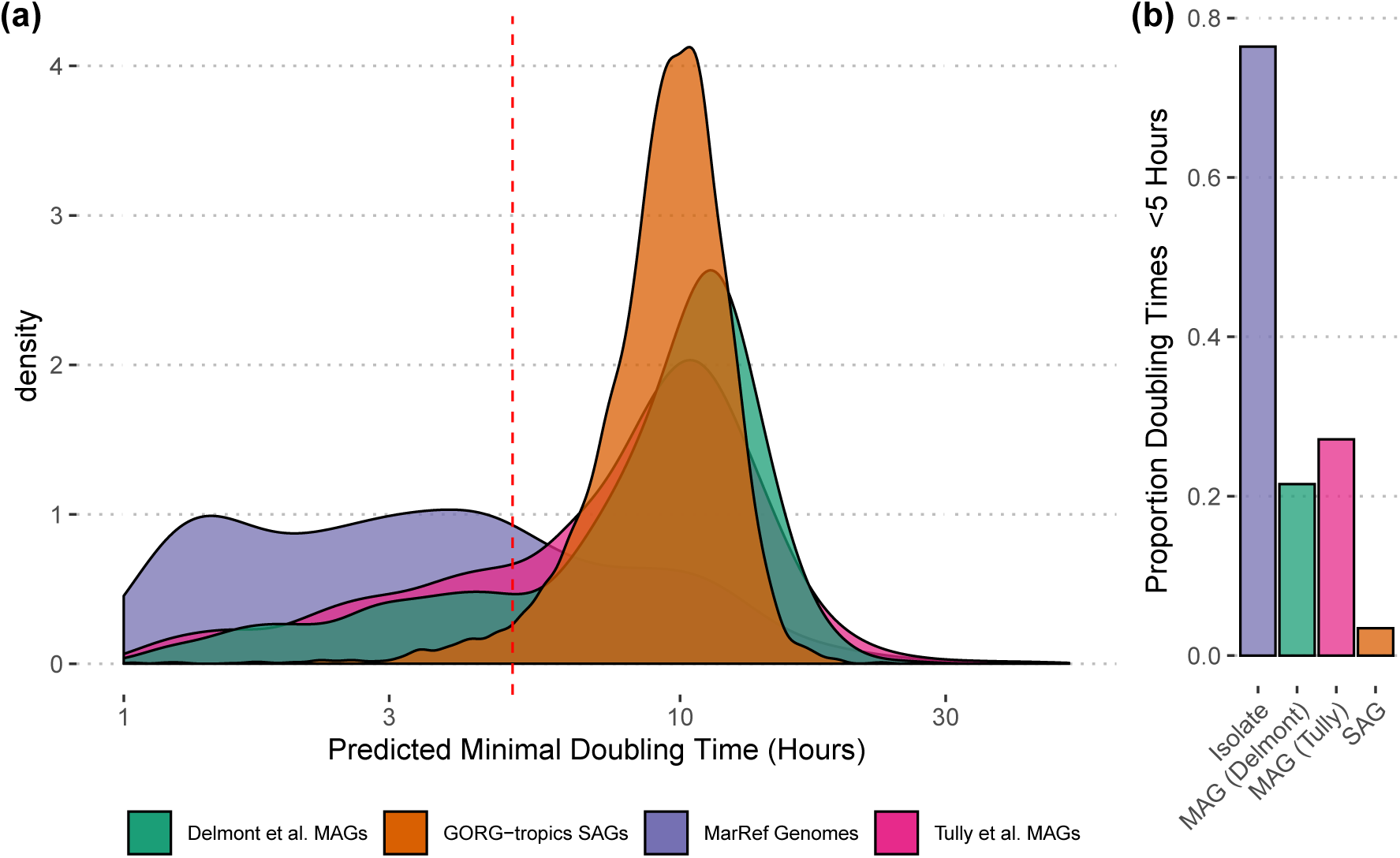
Predicted maximal growth rates in marine environments. Observe that (a-b) genomes from isolates have shorter predicted doubling times on average than MAGs and SAGs, and fail to capture the slow-growing fraction of the community. Additionally, SAGs showed a lower overall growth rate than MAGs, with very few doubling times predicted to be under 5 hours. This may be due in part to how SAGs were sampled (only at the ocean surface, rather than at multiple depths), or to some systematic bias in how MAGs are assembled and binned. MAGs generated by distinct research groups showed surprisingly consistent maximal growth rate distributions. Vertical dashed red line in (a) at 5 hours for reference.

**Figure 5:**
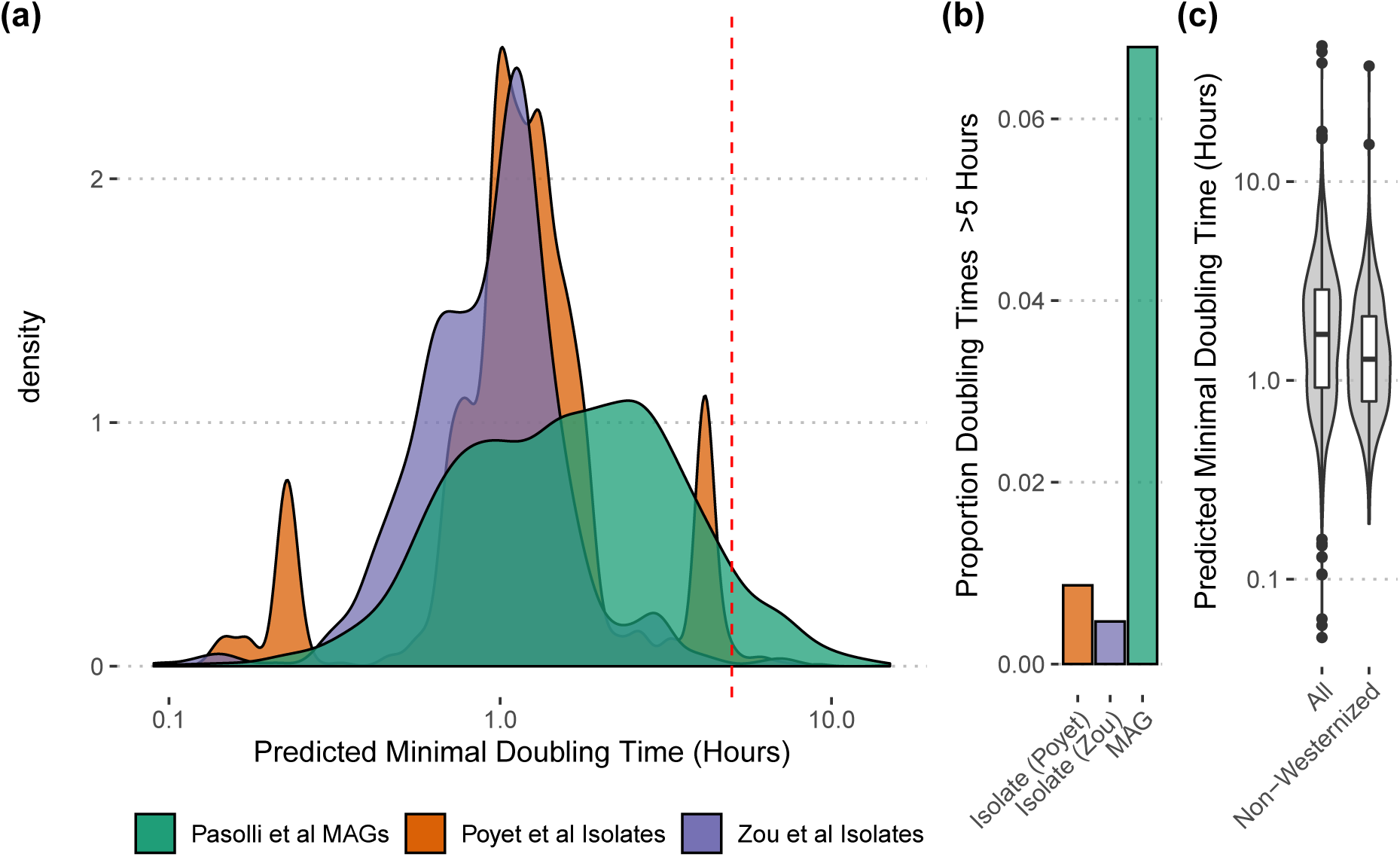
Predicted maximal growth rates in the human gut. Observe that (a-b) genomes from isolates have shorter predicted doubling times on average than MAGs, and fail to capture the slow-growing fraction of the community. Notably, growth-rate distributions are consistent across MAG datasets (S18 Fig) in the gut, though they vary across body sites (S19 Fig). We also found that (c) gut microbes associated with non-westernized microbiomes had slightly higher growth rates than gut microbes in general. Vertical dashed red line in (a) at 5 hours for reference.

Finally, we note that there are many potential use-cases for gRodon and the EGGO database, especially when studying subsets of microbes for which additional metadata is available. For example, microbes associated with “non-westernized” human gut microbiota have a significantly shorter doubling time on average than the gut microbiome as a whole (t-test, *p* = 4.1 *×* 10^*-*6^; Fig 5c; classification of “non-westernized” taxa from [49]; we note that this terminology centers a mythic monolithic “Vest” as a reference against which all other groups are to be compared, and should be revised [32]), perhaps indicating that they are primarily infrequent but fast-growing community members caught during a bloom. As another example, the very largest cells in marine samples seem to also be those with the highest maximal growth rates (Fisher’s exact test, *p* = 2.2 *×* 10^*-*15^; S15 Fig). This is consistent with the “nutrient growth law” coined by Schaechter et al [57], which describes a simple exponential relationship between bacterial cell volumes and their growth rates. Because maximal growth rate is a basic parameter of microbial lifestyle [55], gRodon and EGGO allow us to build better large-scale comparative studies linking specific traits and habitats to particular microbial life-histories.

### Using EGGO to predict growth using only 16S rRNA

There are many organisms for which we do not have genomic information, but for which we have the genomic information of a close relative. Vieira-Silva et al. [72] observed conservation of growth rate below the genus level. We leverage these phylogenetic relationships alongside our comprehensive EGGO database to drastically expand the set of organisms whose growth rates we can predict.

The growth rates of species pairs within a genus are strongly associated. This is true looking at actual maximal growth rates (linear regression, *p* = 2.4 *×* 10^*-*4^, *R*^2^ = 0.39, despite a small number of datapoints *n* = 25), but becomes more apparent when we examine the large number of inferred growth rates in EGGO (linear regression, *p <* 2.2 *×* 10^*-*16^, *R*^2^ = 0.42; S16 Fig). In order to assess how closely two organisms must be related to reliably extrapolate maximal growth rate, we built a phylogeny of 16S rRNA sequences with corresponding records in EGGO. We predicted maximal growth rate as the weighted geometric mean of an organism’s nearest 5 relatives on the tree (weighted by inverse patristic distance, see Methods). Comparing an organism’s entry in EGGO to values extrapolated from closely related relatives, we found that the two quantities were highly correlated (Pearson correlation of log-transformed doubling times *ρ* = 0.78, *p <* 2.2 *×* 10^*-*16^; Fig 6a). Prediction error was relatively insensitive to how distant these neighbors were up to a patristic distance of *∼* 0.2 (Fig 6b; consistent with previous observations [72]). We obtained similar results when predicting only on the basis of the closest relative (Pearson correlation of log-transformed doubling times *ρ* = 0.60, *p <* 2.2 × 10^*-*16^; S17 Fig). Importantly, prediction using a 16S tree relies on a large database of pre-predicted maximal growth rates (i.e., EGGO), meaning that errors are compounded over multiple rounds of prediction. We thus caution against over-interpretation of phylogenetic predictions, though these predictions can offer a useful baseline estimate for organisms for which we have very little life-history information. One option for the conservative microbiologist is to use phylogeny to predict whether an organism is a copiotroph or oligotroph (following our earlier cutoff of a 5 hour doubling time), as classification is generally an easier task than regression. Our approach to phylogeny-based prediction did well when applied for classification of oligotrophs (i.e., whether an organism had a doubling time *>* 5 hours; accuracy= 0.98, Cohen’s *κ* = 0.61). We include a blast database of 16S sequences for organisms with records in EGGO alongside the online database so that users may search their own 16S sequences to predict growth.

**Figure 6:**
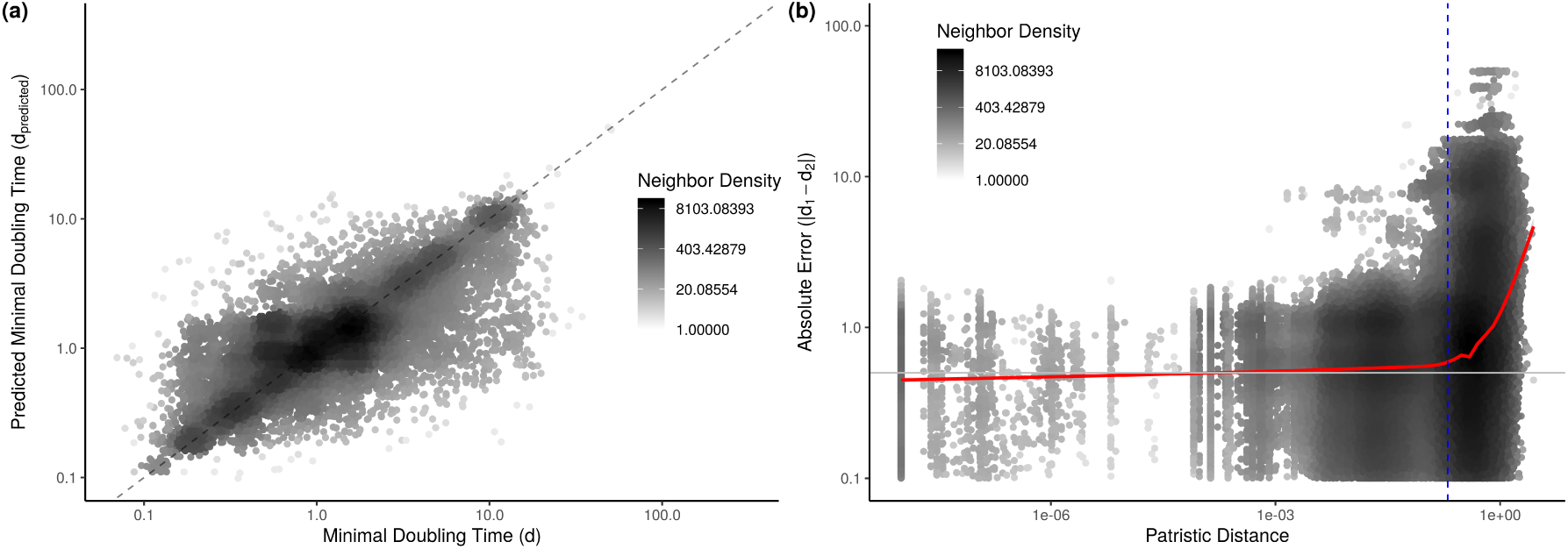
Closely related organisms have similar predicted maximal growth rates. (a) We predicted the growth rate of an organism based on closely related organisms in EGGO and found good correspondence to that organism’s entry in EGGO. Dashed line denotes the *x* = *y* line. (b) Pairs of randomly sampled organisms have similar growth rate entries in EGGO as long as they are closely related (vertical dashed blue line at a patristic distance of 0.2, the same threshold found in [72]). Horizontal gray line at *d* = 0.5 hours for reference. (a-b) Points shaded relative to number of nearby neighbors in order to visualize density (ggpointdensity R package https://github.com/LKremer/ggpointdensity).

## Conclusions

We produced a community resource in the form of an easy-to-use and well documented R package (gRodon) and comprehensive database (EGGO) for predicting and compiling maximal growth rates across prokaryotes. Using these tools we show how existing cultured isolates do not fully capture the diversity of prokaryotic lifestyles. We are unlikely to overcome these biases easily, as the slow-growing microbes missing from our culture collections are precisely the ones we expect to be most difficult (and time-consuming) to grow in the laboratory. Yet, we have their genomes, and may be able to extrapolate their traits from microbes that are more easily cultivable. Growth rate is one example where inference of traits from genomes has clear utility, though we emphasize that genome-wide signals may be confounded by other evolutionary and/or demographic processes and that it is important to assess their robustness and limitations, as we have done here.

Finally, we emphasize that the relationship of the in situ growth rate and the maximal growth rate of an organism is not clear given the cryptic influence of top-down and bottom-up controls at the sampling time. There are any number of reasons why an organism may not reproduce at its physiological maximal rate (e.g., fluctuating habitat quality, dispersal to sub-optimal habitats, etc.). Nevertheless, it is encouraging that recent work using natural communities has shown that CUB-based estimators do a reasonably good job of predicting observed instantaneous growth rates in marine systems [36], even as peak-to-trough [30, 4, 19, 17] methods of estimating growth have been reported to work poorly for marine plankton, with the exception of the most highly abundant copiotrophs [36]. Thus, taken together with our benchmarking against nutrient-enrichment experiments, the data suggest that CUB-based estimators of maximal growth rate tend to also recapitulate the instantaneous growth rate of a community.

## Methods

All scripts used to generate figures and analysis, as well as predicted growth rates for various genomic datasets and the full EGGO database, are available at https://github.com/jlw-ecoevo/eggo. The gRodon package, including documentation and a vignette can be downloaded at https://github.com/jlw-ecoevo/gRodon.

### Model Fitting

For each species with a growth rate listed in the original Vieira-Silva dataset (214; [72]) we downloaded all available complete genome assemblies from NCBI’s RefSeq database (1415; [65, 66, 23]). For each species we calculated the mean of each of our three codon usage statistics across all genomes corresponding to that species. Ribosomal protein annotations were taken directly from the annotations generated by NCBI’s default prokaryotic annotation pipeline, and these were the ribosomes passed to both growthpred and gRodon. Importantly, growthpred can also search for ribosomal proteins using a provided database, though we did not use this feature so as to make sure the two prediction methods were compared on identical datasets. For initial model fitting, we excluded thermophiles and psychrophiles from the dataset (31) as these organisms systematically differ in their codon usage patterns [72]. Similar to growthpred, we include a temperature option fit using these microbes in the final gRodon package that accounts for optimal growth temperature in the final model, though given the few extremophiles used to fit this model we caution users against drawing strong conclusions when it is applied to extremophiles (by default temperature is not used for prediction).

We then fit a linear model to box-cox transformed doubling times (optimal *λ* chosen using the MASS package [71]) using our three codon usage measures as predictors. Similarly we fit models for gRodon’s “partial” (excluding pair-bias) and “metagenome” (excluding pair-bias and consistency) modes.

For fitting on the Madin et al. [38] training set we used the same model fitting procedure. We took the minimal recorded doubling time from each species in the “condensed_traits_NCBI.csv” supplementary file (https://doi.org/10.6084/m9.figshare.c.4843290.v1), and where possible obtained all completely assembled genomes associated with that species from RefSeq. This yielded our training set with 464 species matched to 8287 genomes. Notably, 130 of these species were either thermophiles or psychrophiles, perhaps making this training set preferable when dealing with extremophiles.

The Gaussian-mixture model in Fig 3 was fit using the Mclust() function in the mclust package with default settings [58]. Mclust chooses the optimal mixture of Gaussian based on BIC and finds this optimum (for mean and variance) using an expectation-maximization algorithm.

### Metagenomic Data

The raw sequencing data for the metagenomic water samples taken at the end of the Okie et al. [45] experiments were obtained from NCBI under BioProject PRJEB22811. Raw sequencing data for the time-series samples taken by Coella-Camba et al. [9] were obtained from NCBI under BioProject PRJNA395437. Adapters and low quality reads were trimmed using fastp v0.21.0 [7] with default parameters and only reads longer than 30 bp were kept for further analysis. Okie et al. [45] samples were assembled individually using metaSPAdes v3.10.1 [43]. Coello-Camba et al. [9] samples were assembled individually using megahit v1.2.9 [34] with default parameters. We called and annotated ORFs from assemblies using prokka [59] (with options “--metagenome --compliant -- fast”). Reads were mapped to ORFs using bwa 0.7.12 [35], and the number of reads aligned to each ORF were counted using bamcov v0.1.1 (available at https://github.com/fbreitwieser/bamcov). We ran gRodon in weighted and unweighted metagenome mode on each sample, with weights corresponding to mean coverage depth (corrected for gene length). In weighted metagenome mode the median CUB of the highly expressed genes is taken as a weighed median (weightedMedian in matrixStats R package), with weights corresponding to mean depth of coverage for that gene. One sample from Coella-Camba et al. [9] had a very atypical estimated average minimal doubling time over twice as long as any other estimated doubling time from this dataset (MG078 at 3.1 hours, as compared to the second longest doubling time in MG002 at 1.4 hours), and strongly disagreeing with a replicate sample from the same experiment and timepoint (MG073 at 0.35 hours). Upon closer inspection, this sample had far fewer bases than the rest (133M bases vs *>* 1G bases) and only a little over 400 genes were detected in the assembly, far too few for accurate assessment of community-wide growth rate, leading us to omit this sample from further analyses.

### EGGO Datasets

We downloaded all prokaryotic assemblies from RefSeq [65, 66], as well as several collections of isolate genomes [29, 50, 75], MAGs [69, 49, 41], and SAGs [46]. Where possible, we used per-existing gene annotations provided by NCBI. For the Pasolli et al. [49] and Nayfach et al. [41] MAGs gene predictions were not available and we used prokka to predict ORFs and annotate ribosomal proteins [59]. Note that for both of these MAG datasets we used a subset of all MAGs designated as being representatives of species clusters by the authors. We then ran gRodon on each genome, using partial mode for MAGs and SAGs (which vary in their completeness). Finally, we filtered results from genomes with few ribosomal proteins. Similar to Vieira-Silva et al [72], we found that growth rates were biased when <10 highly expressed genes were used for prediction (S20 Fig), and we used this cutoff for our MAGs and SAGs. For our isolate genomes this generally was not an issue, with over 99% of genomes in RefSeq having between 50-70 annotated ribosomal proteins. We filtered all genomes outside this range to remove a very small set of obvious problem cases (e.g., one Bacillus genome that had over 1000 annotated ribosomal proteins). The numbers in Table 1 correspond to post-filtering genome counts.

### Measuring Bias

We use the MILC measure of codon usage bias [63] implemented in the coRdon R package [16]. This bias measure behaves slightly better than the ENC’ measure used by Vieira-Silva et al [42, 72], and automatically accounts for the CUB of genomic background in its calculation (by taking the genome-wide distribution of codons as its expected distribution; [63, 16]). As recommended in the coRdon documentation, genes with fewer than 80 codons were omitted from our calculations. Importantly, we calculate the MILC statistic on a per-gene basis rather than concatenating all of our genes. The contribution (*M*_*a*_) of each amino acid (*a*) to the MILC statistic for a gene is calculated as:

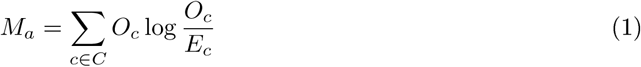

where *C* is the set of codons coding for *a, O*_*c*_ is the observed count of codon *c*, and *E*_*c*_ is the expected count of codon *c* (See the original paper for the full calculation of the MILC statistic; [63]). Typically, *E*_*c*_ for a given gene is estimated using the genome-wide frequency of that codon *c*. This is what we mean when we say that for our CUB measurement the bias of highly expressed genes is calculated “relative to the genomic background”.

For our consistency calculation MILC was also used, but was calculated using the highly expressed proteins as the expected background (using the “subset” option in coRdon). In other words, we estimated the expected count of a codon, *E*_*c*_, using the frequency of that codon in highly-expressed genes only, rather than the genome-wide frequency.

For codon-pair bias we implemented the calculation by Coleman et al. [10] that controls for background amino acid and codon usage when estimating the over/under representation of codon pairs (see their S1 Fig for relevant equation).

### Population Parameters

We obtained estimates of *N*_*e*_ from [3], which are based on dN/dS ratios (the intuition being that selection acts more efficiently in large populations). Gene-specific recombination rates were obtained by applying the PhiPack [5] program for detecting recombination to the ATGC database of closely-related genome clusters [31], as described in Weissman et al. [73].

### Extrapolating Between Closely Related Taxa

For all genomes used to build EGGO we extracted all annotated 16S rRNA genes and then aligned these sequences and removed poorly-aligned columns using ssu-align and ssu-mask (default settings; [40]). We then filtered sequences for which less than 80% of positions were accounted for (i.e., were gaps). We ran fasttree on the resulting alignment (with -fastest, -nt, and -gtr options; [51]) to obtain a phylogeny with 192,195 tips representing 60,421 organisms. For phylogenetic prediction of maximal growth rate we then omitted any tips with EGGO entries where *d >* 100 hours (13 tips) to minimize the influence of outliers.

To predict growth rate we first randomly sampled one tip per organism in our tree (to avoid predicting an organisms growth rate from itself). We then iteratively found the five closest tips to each tip in the tree, and took the weighted geometric mean of the growth rates associated with these tips. This gave us our predicted maximal growth rate on the basis of 16S rRNA in Fig 6a. Weights were calculated as inverse patristic distance, with a small constant added for when organisms had identical 16S sequences (e.g., multiple genomes in EGGO for the same species):

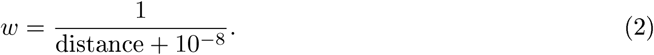

For S17 Fig, the predicted rate was simply taken as the rate associated with the closest tip on the tree. We identified the closest tips using the castor R package [37].

To produce Fig 6b we sampled 10,000 tips from our tree and calculated all pairwise distances between tips using the cophenetic.phylo() function in the ape R package [47].

## Acknowledgments

JLV was supported by a postdoctoral fellowship in marine microbial ecology from the Simons Foundation (award #653212). We also acknowledge support from the Simons Foundation Collaboration on Computational Biogeochemical Modeling of Marine Ecosystems/CBIOMES grant 549943 to JAF, and U.S. National Science Foundation grant OCE 1737409 to JAF.

## Notes

https://github.com/jlw-ecoevo/eggo

https://github.com/jlw-ecoevo/gRodon

### Competing Interest Statement

The authors have declared no competing interest.

