## Supplemental Figures for "Estimating maximal microbial growth rates from cultures, metagenomes, and single cells via codon usage patterns"

July 25, 2020

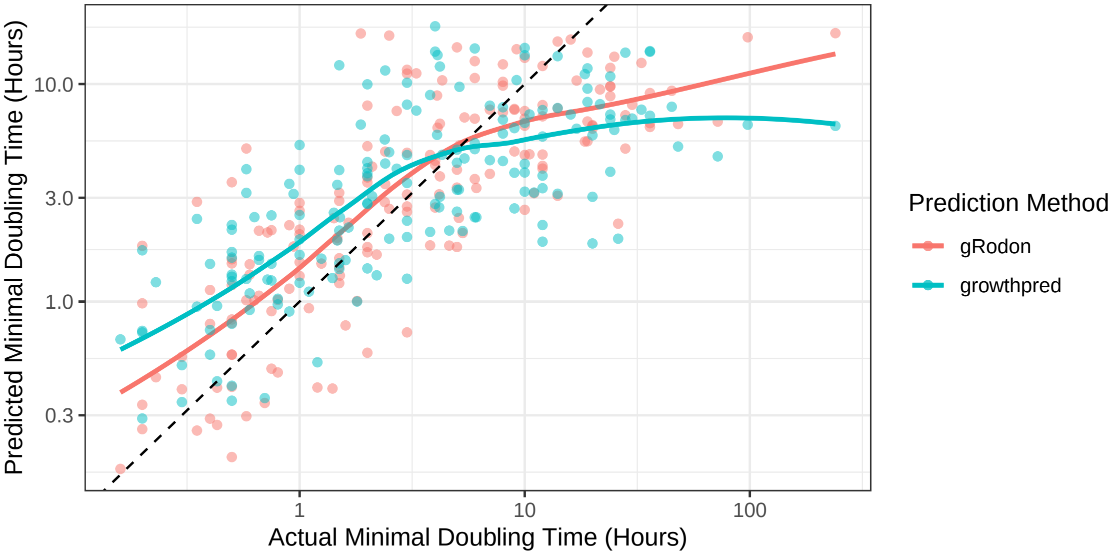


Figure S1: gRodon fits the data better than growthpred at both long and short doubling times.

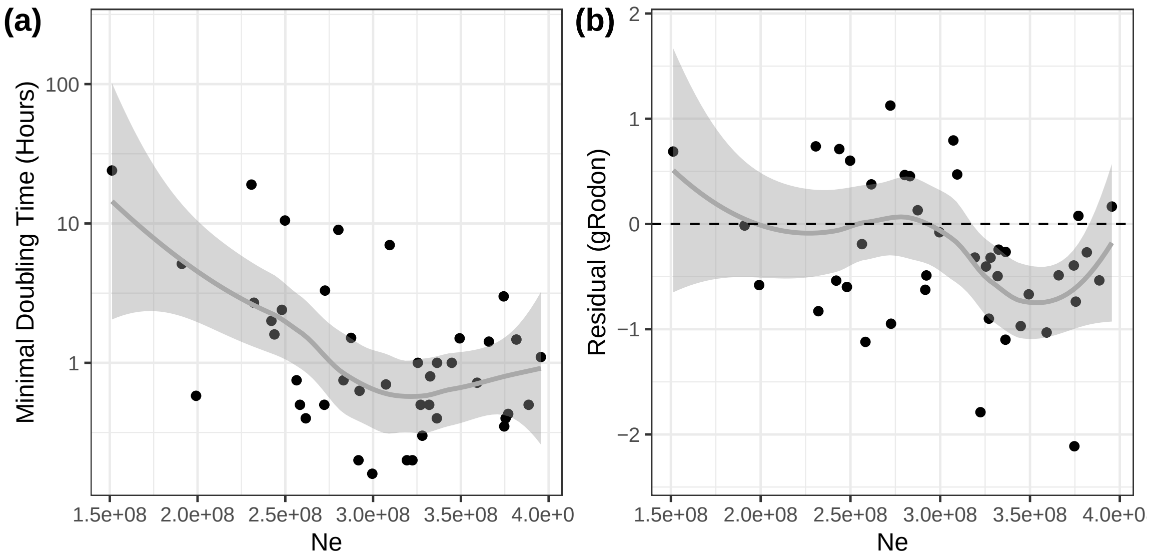


Figure S2: Effective population size is associated with growth rate ($p = 3.2 \times 10^{-4}$, $R^2 = 0.26$, linear regression) and has a minor effect on our model residuals ($p = 0.018$, $R^2 = 0.11$, linear regression).

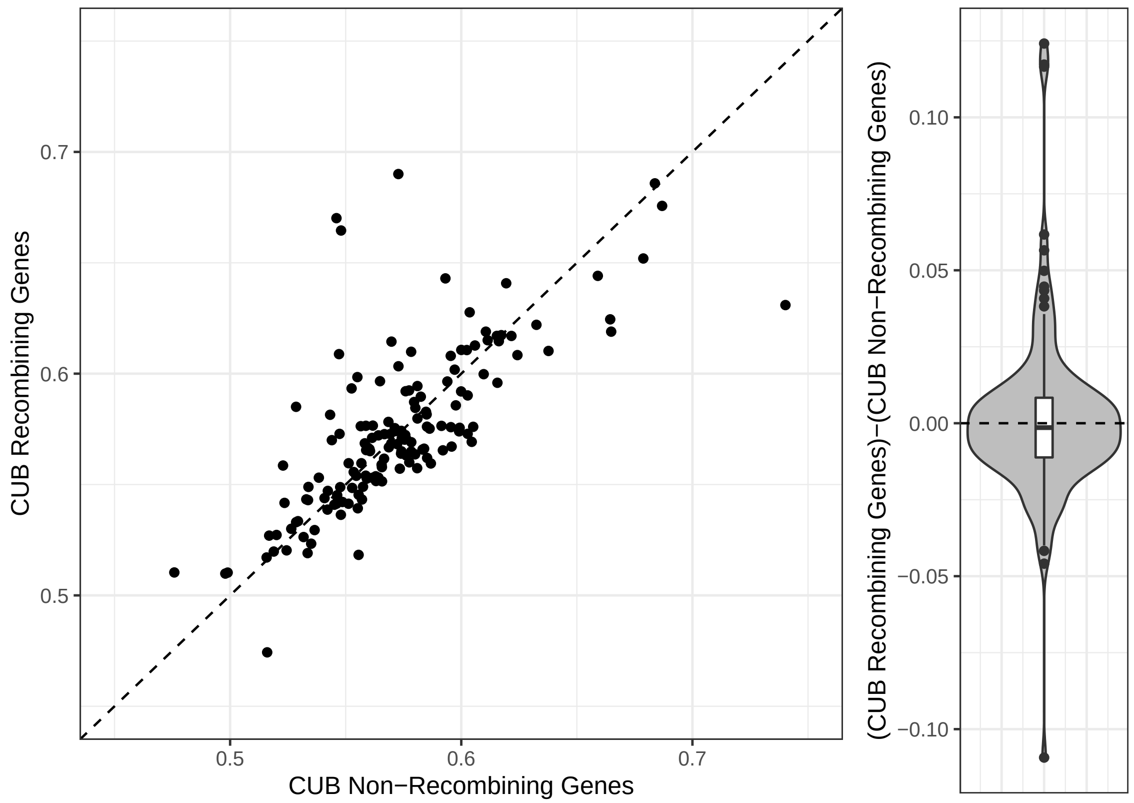


Figure S3: Recombining genes do not have higher CUB than non-recombining genes. Each point represents a cluster of closely related genomes.

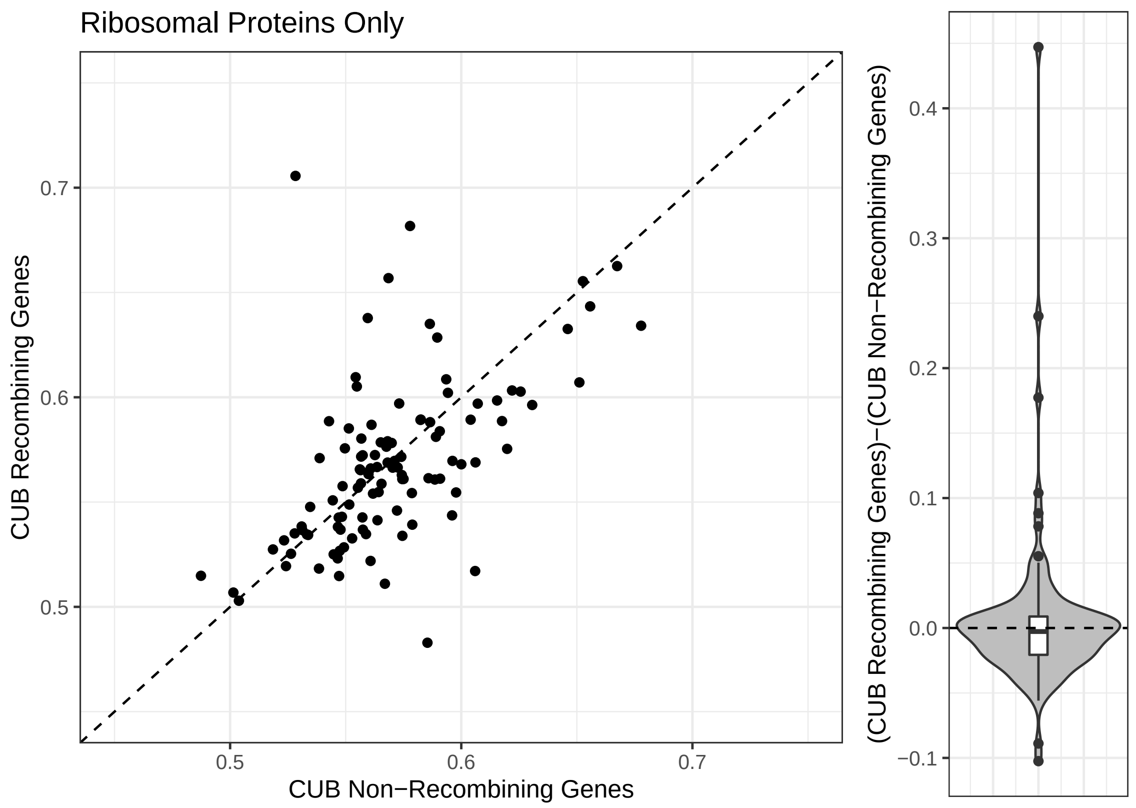


Figure S4: Recombining genes do not have higher CUB than non-recombining genes, even when restricting our analysis to only genes coding for ribosomal proteins. Each point represents a cluster of closely related genomes.

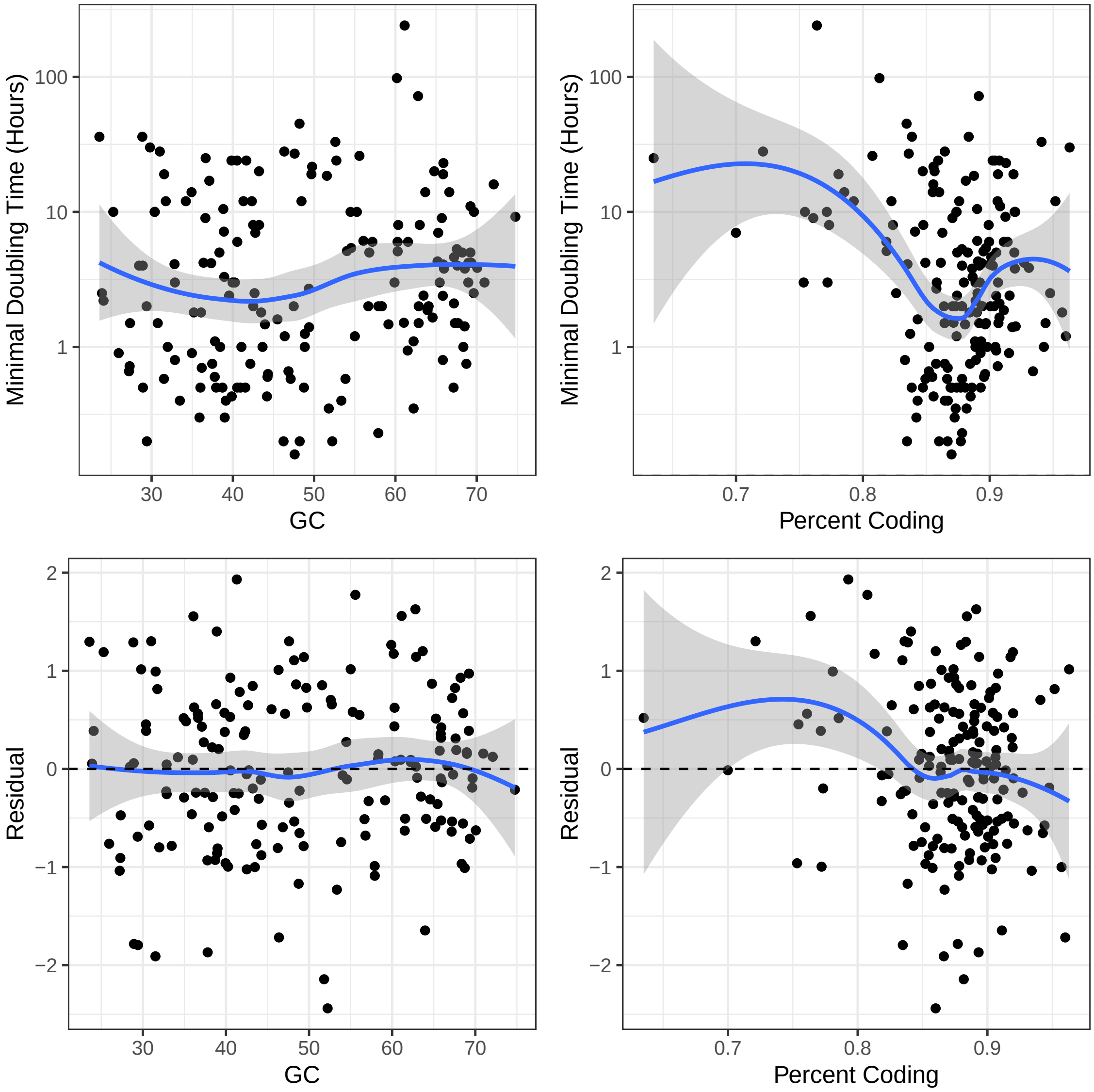


Figure S5: Genomic GC content and percent coding sequence have no effect on gRodon model residuals.

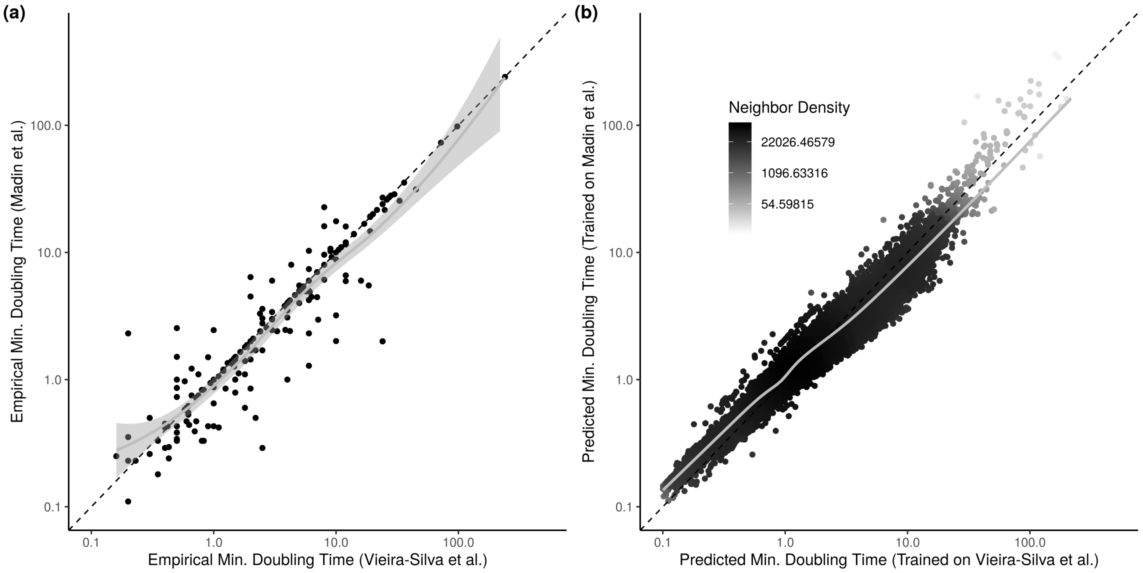


Figure S6: While (a) the default and alternative training sets disagree on the minimal doubling times of a number of species, they are overall highly correlated (Spearman correlation, $\rho = 0.98$, $p < 2.2 \times 10^{-16}$). (b) The predictions of the models trained on these datasets (complete EGGO database; 217,074 predictions) are even more tightly correlated (Spearman correlation, $\rho = 0.9997$, $p < 2.2 \times 10^{-16}$). Points shaded relative to number of nearby neighbors in order to visualize density (ggpointdensity R package).

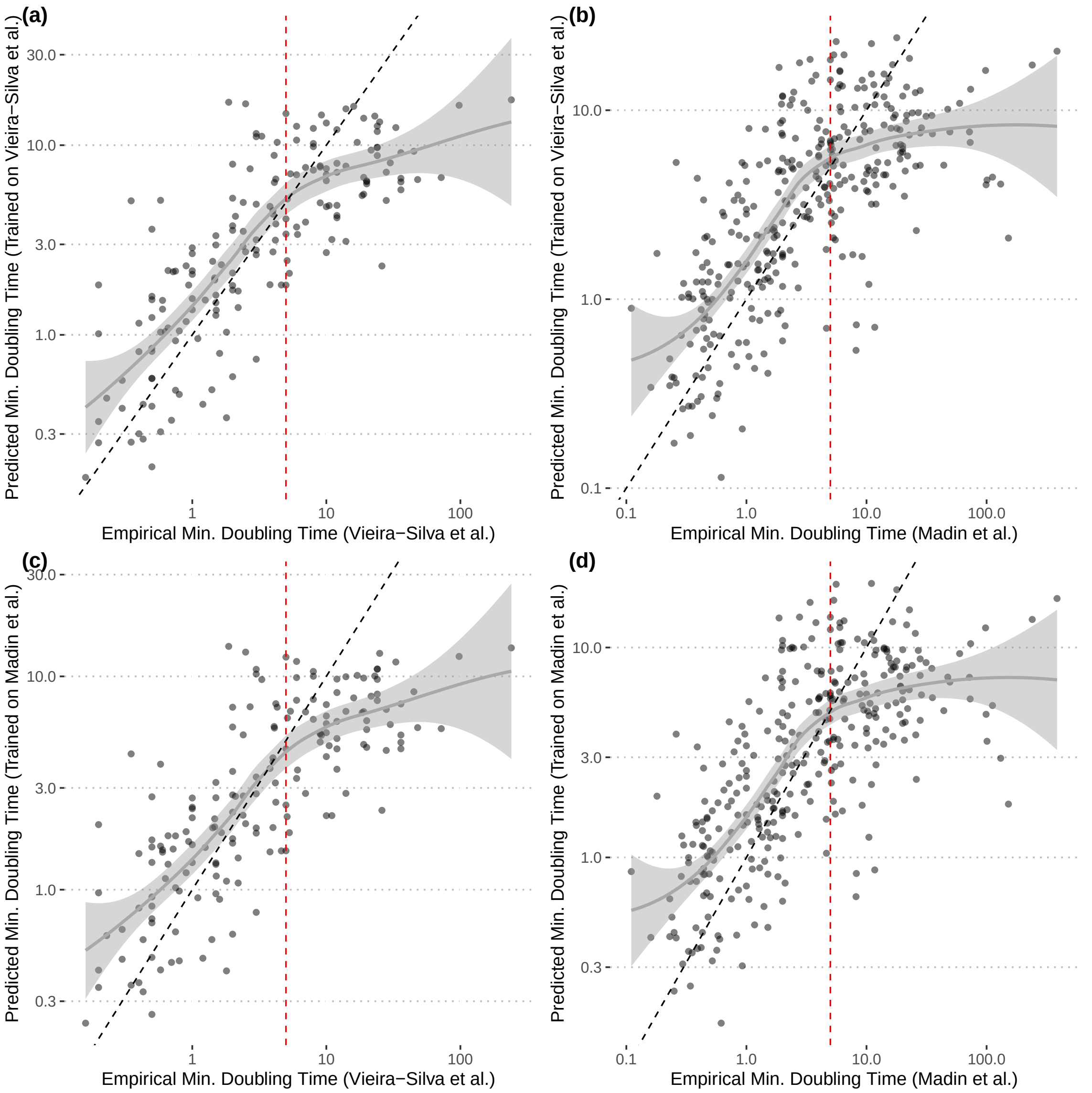


Figure S7: Model fit is largely insensitive to training data. (a-b) Model trained on default training set. (c-d) Model trained on alternative training set. Notice that fits of both models on either set appear very similar (a vs. c and b vs. d).

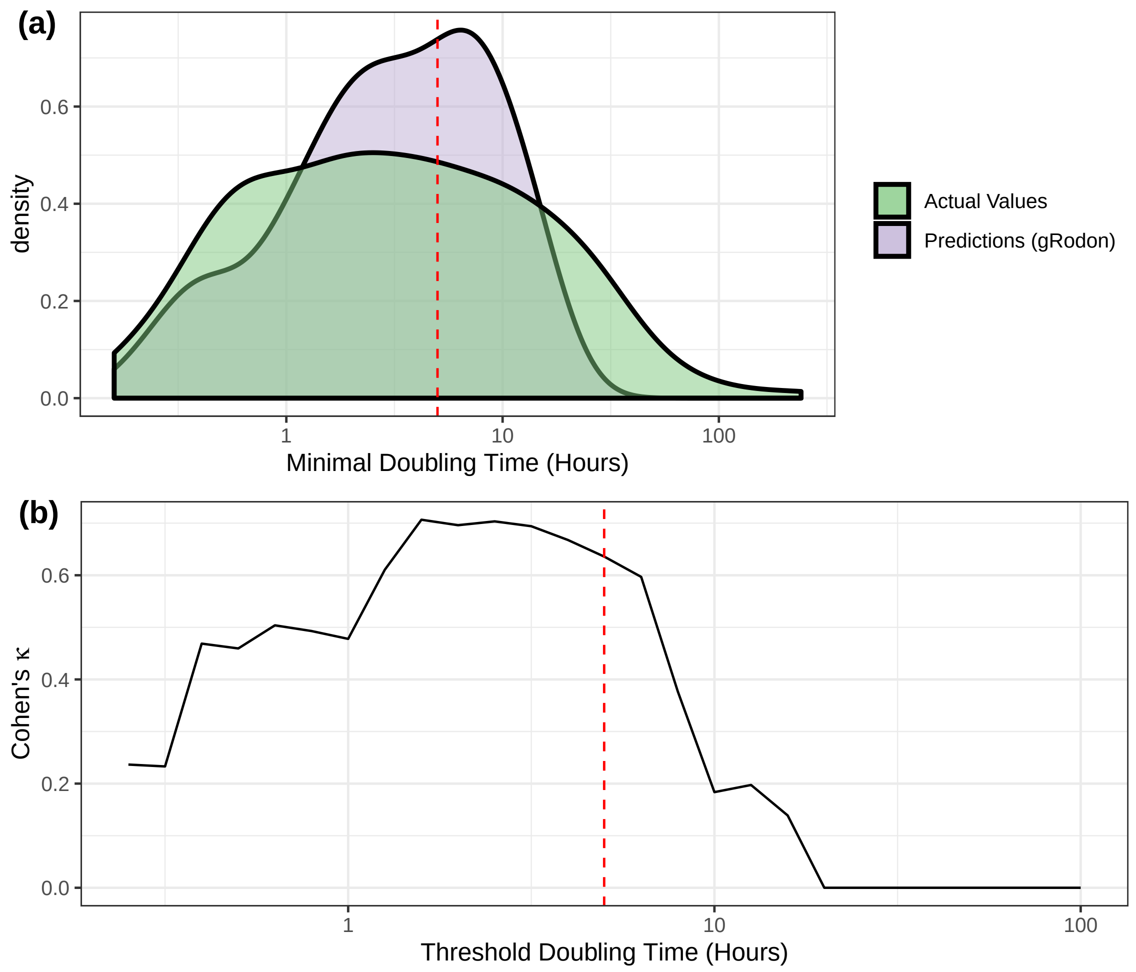


Figure S8: For long doubling times, 5 hours is an appropriate threshold to classify oligotrophs. (a) Above 5 hrs, doubling times are underestimated by gRodon, leading to a bimodal distribution of predicted growth rates. (b) Varying the threshold for classifying fast/slow growers, we see a steep drop-off in performance above 5 hrs. Cohen's $\kappa$ is a measure of performance that corrects for unbalanced classes (values above zero indicate predictive ability above a null model that always guesses the majority class).

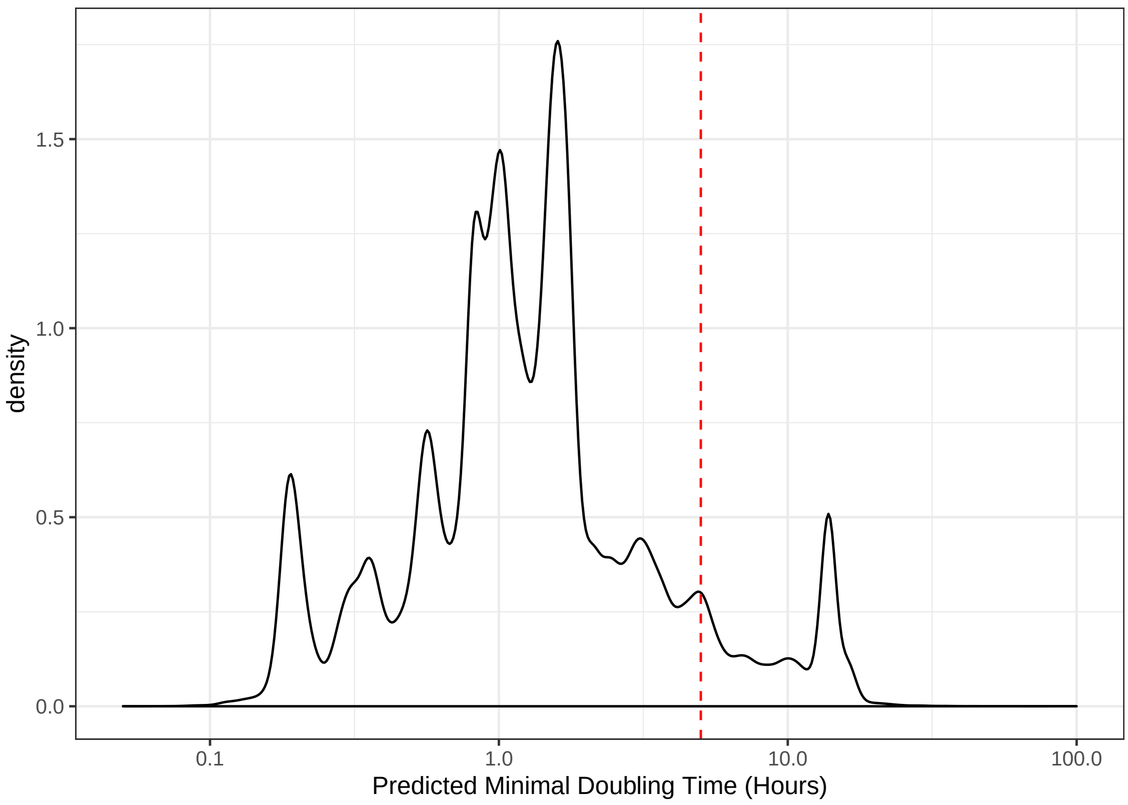


Figure S9: Distribution of all growth rates predicted from RefSeq genomes. Uneven sampling of taxa leads to several irregular and sharp peaks.

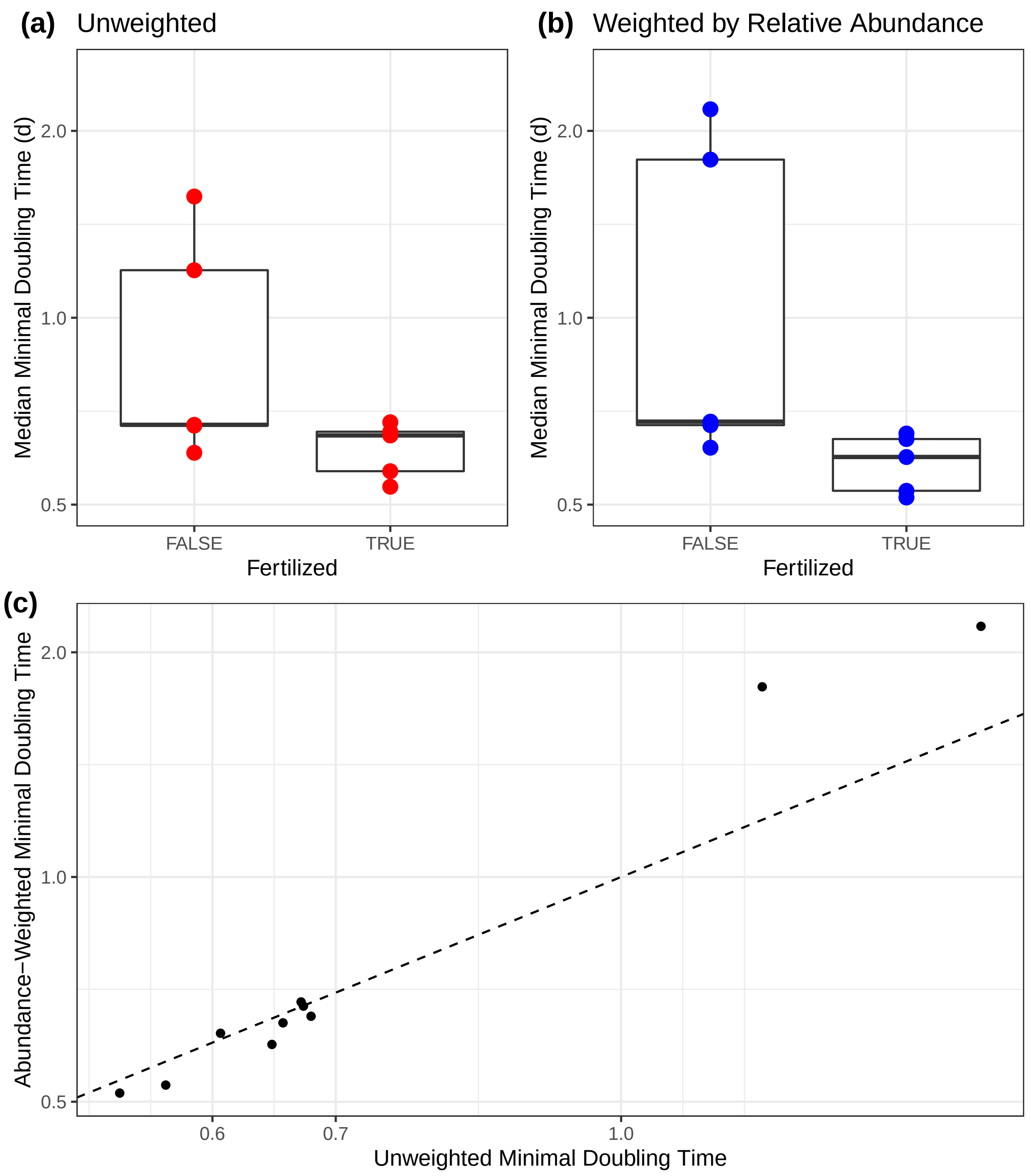


Figure S10: Average community-wide maximal growth rate predictions from a nutrient enrichment experiment. Note that (a) gRodon's unweighted metagenome mode shows less differentiation between treatments than (b) gRodon's weighted metagenome mode, though (c) these two modes have correlated results.

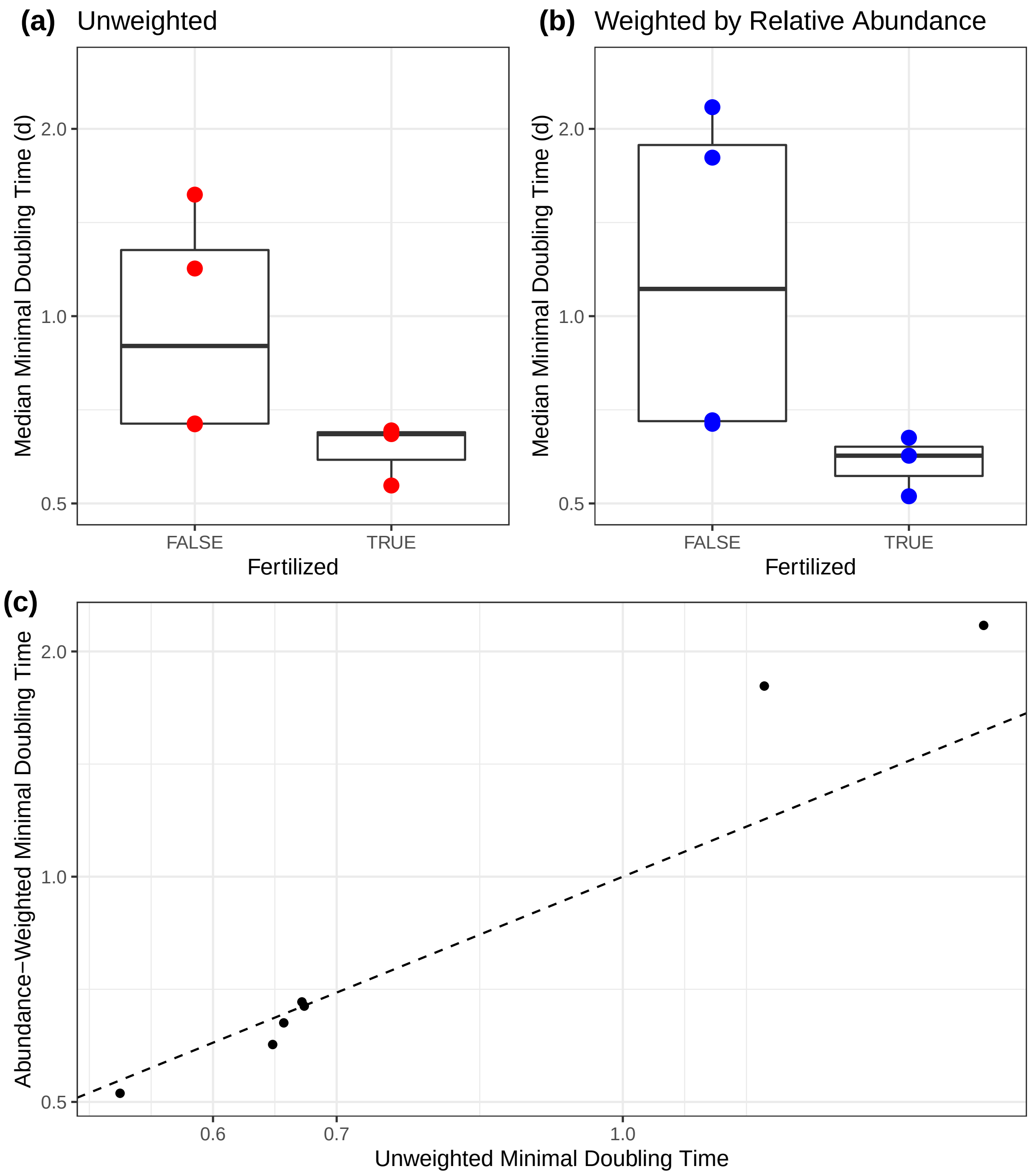


Figure S11: Average community-wide maximal growth rate predictions from a nutrient enrichment experiment. Only includes samples analyzed in the original paper ($n = 7$). See S8 Fig for the full set of samples ($n = 10$). Observe that in (b) weighted metagenome mode all fertilized treatments have shorter average community-wide minimal doubling times than all unfertilized treatments.

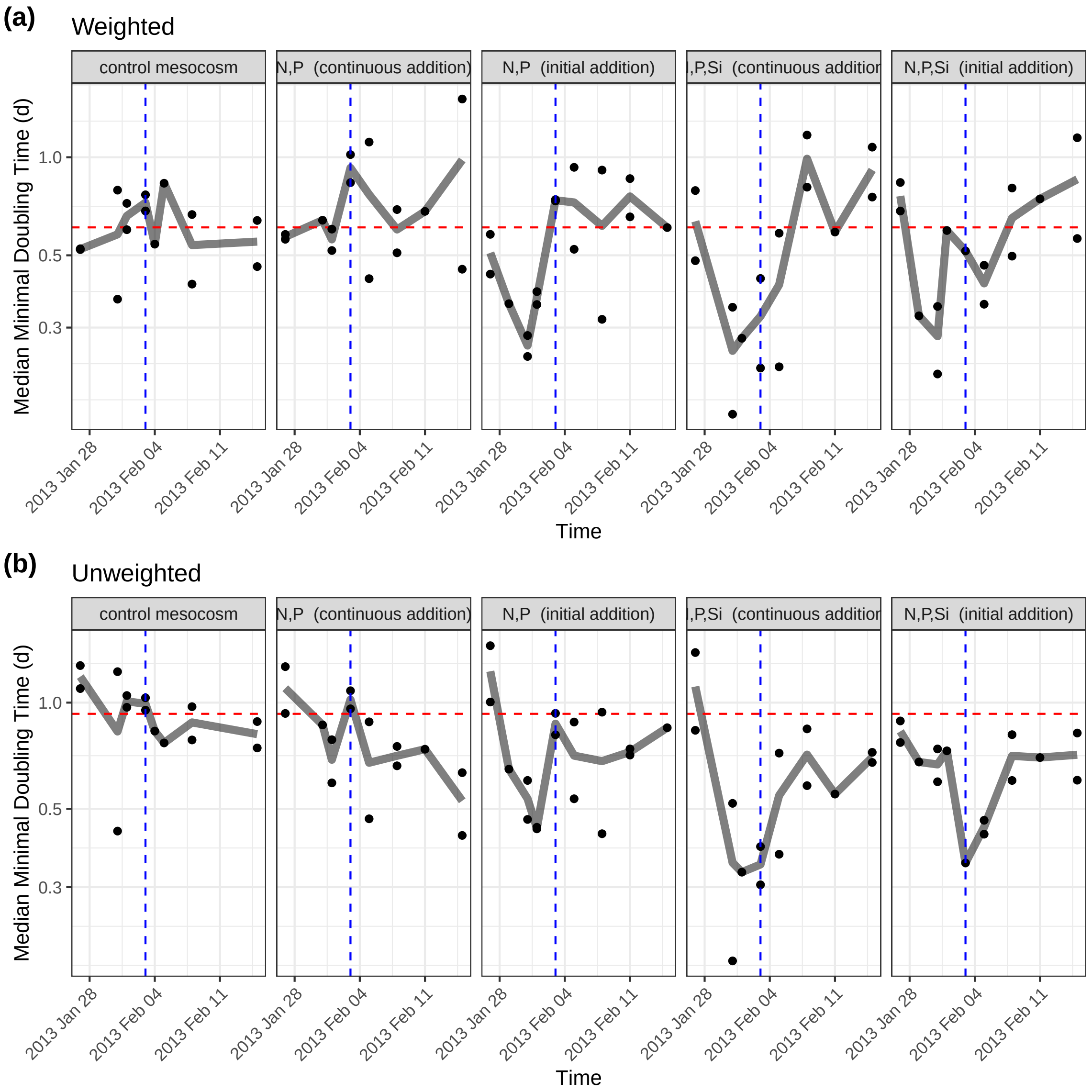


Figure S12: Average community-wide maximal growth rate predictions from a time-series nutrient enrichment experiment in the Red Sea. Coello-Camba et al. added either N+P or N+P+Si to marine mesocosms in either a large initial dose or smaller daily doses. The authors observed large *Synechococcus* blooms in both N+P+Si treatments as well as a smaller bloom in the N+P initial addition treatment within the first week of treatment (blue dashed vertical line), followed by a return to baseline. Both weighted and unweighted metagenome modes detected these changes in the community in these treatments as a steep initial drop in average community-wide minimal doubling time, followed by a return to the initial baseline (red dashed line denotes mean of the control treatment). The control and N+P continuous treatments had very small early *Synechococcus* blooms which gRodon was unable to detect.

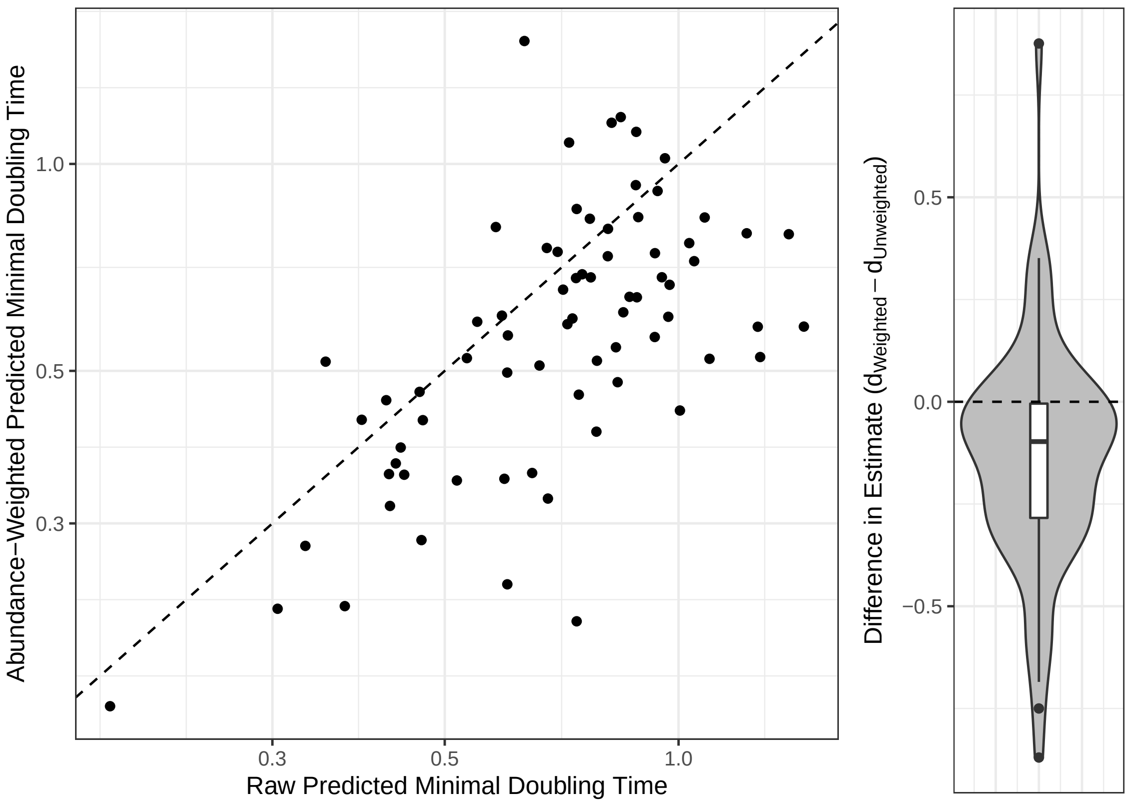


Figure S13: Average community-wide maximal growth rate predictions from a time-series nutrient enrichment experiment in the Red Sea. Coello-Camba et al. added either N+P or N+P+Si to marine mesocosms in either a large initial dose or smaller daily doses. Unweighted metagenome mode systematically predicts longer average community-wide doubling times for samples from this experiment than weighted metagenome mode (paired $t$-test, $p = 4.9 \times 10^{-5}$).

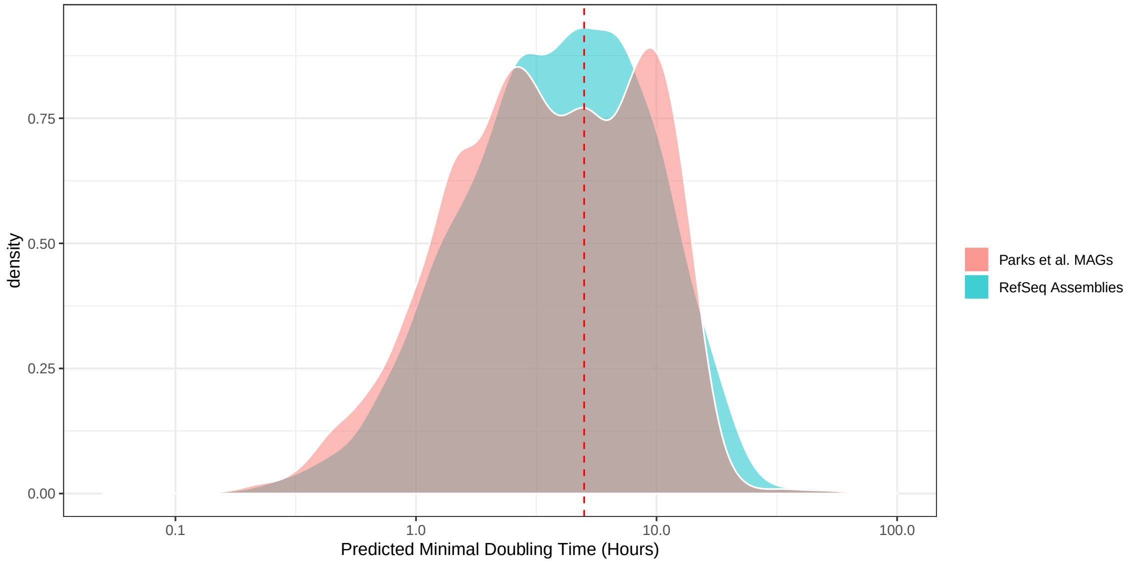


Figure S14: Comparison of growth rates inferred from RefSeq assemblies to those inferred from a large set of MAGs assembled from diverse habitats.

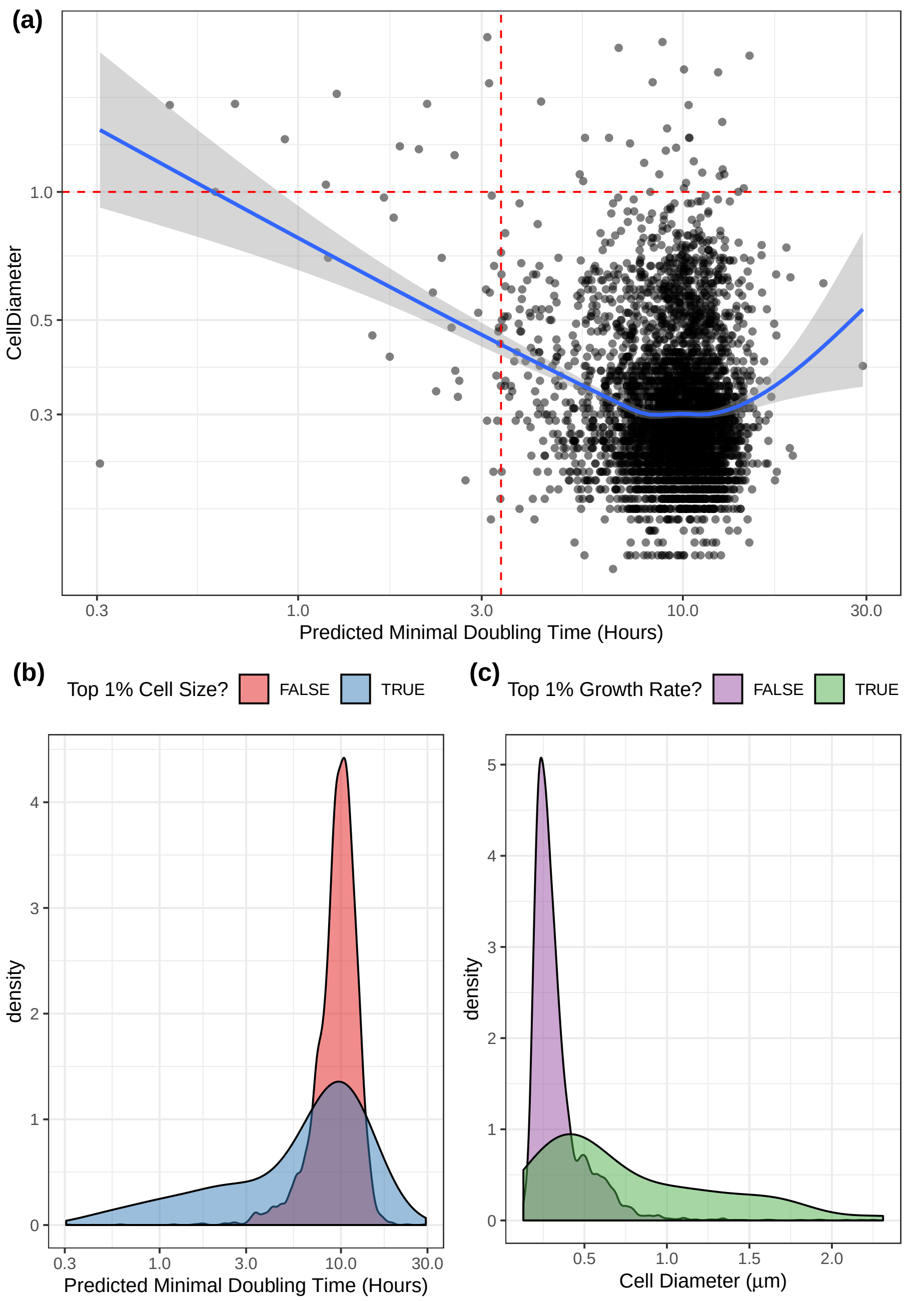


Figure S15: While most cells from the marine surface are small and slow-growing, the biggest %1 of cells also tend to be the fastest-growing 1% (Fisher's exact test, $p = 2.2 \times 10^{-15}$). Cell size measurements were provided in the original GORG-tropics SAG database by Pachiadaki et al.

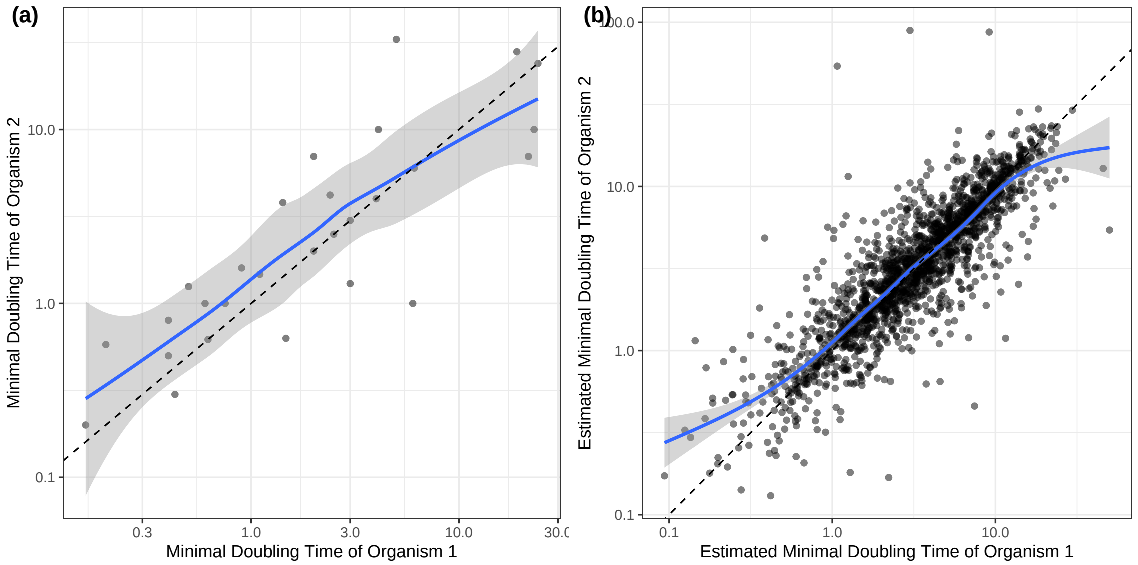


Figure S16: Pairs of organisms in the same genus tend to have similar maximal growth rates. (a) Actual maximal growth rate data for genera in our original dataset (used for fitting gRodon). (b) Inferred maximal growth rates for genera in EGGO (just looking at RefSeq assemblies). In both (a-b) a single pair of species was randomly selected from each genus.

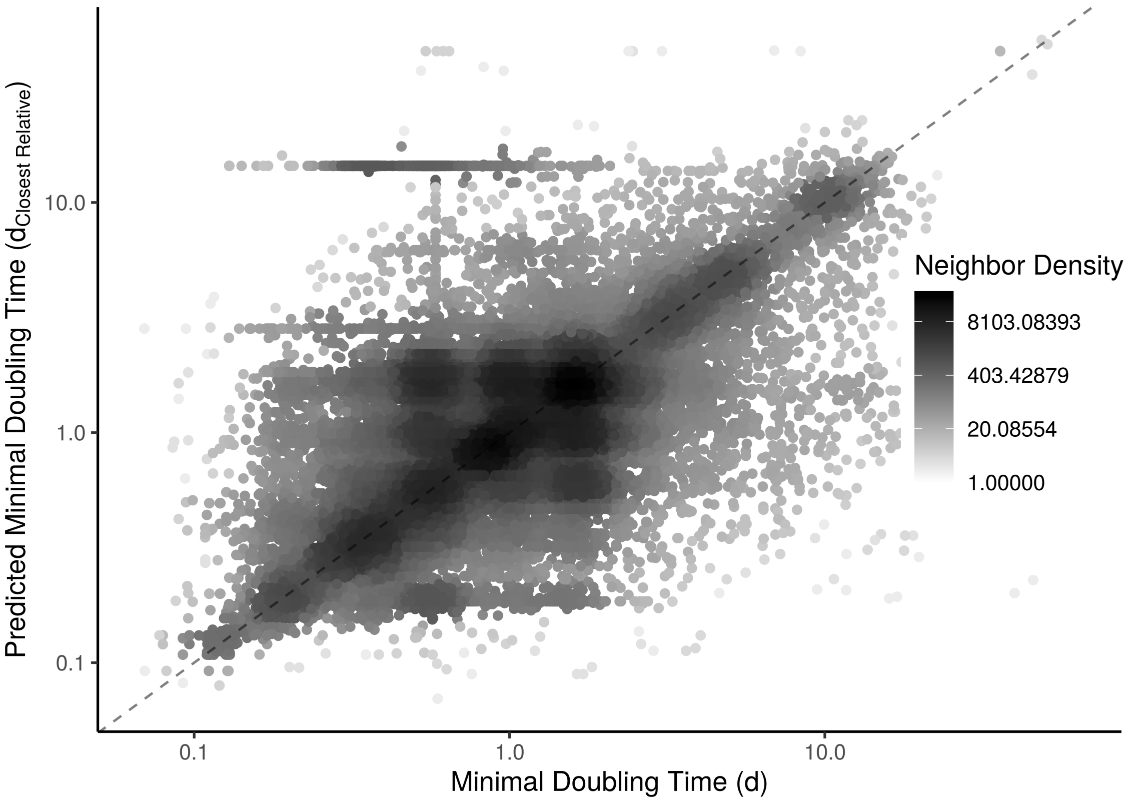


Figure S17: Closely related organisms have similar predicted maximal growth rates. We predicted the growth rate of an organism based on it's most closely related relative in EGGO and found good correspondence to that organism's entry in EGGO. Dashed line denotes the $x = y$ line.

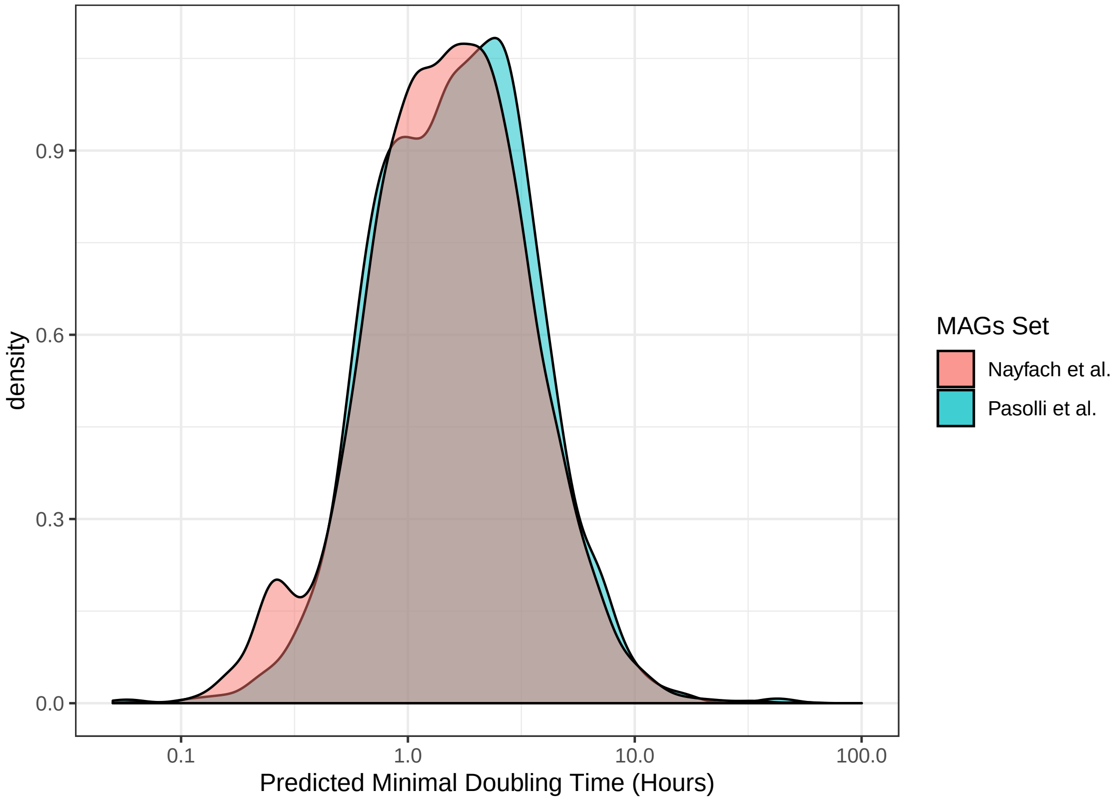


Figure S18: Predicted growth rates from human-gut MAGs are largely consistent across datasets.

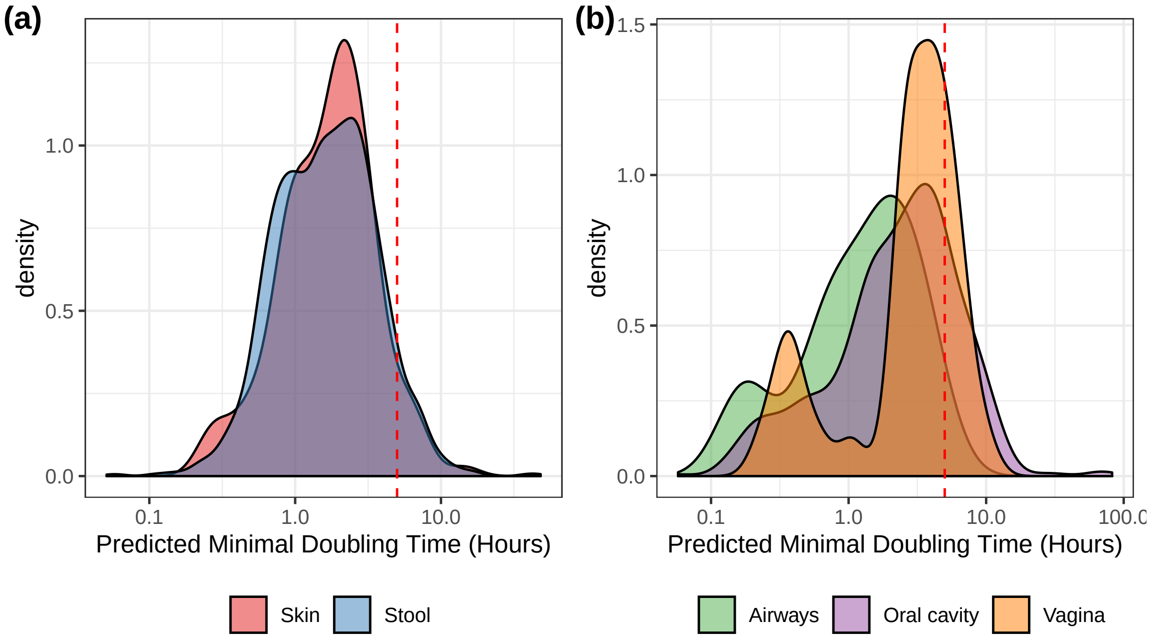


Figure S19: Predicted growth rates from human-microbiome MAGs vary across body sites, with some sites showing a bimodal-distribution of growth rates (the peak at higher doubling times may be our "oligotrophy" peak discussed in the main text, shifted towards doubling times $< 5$ hours because of lower $N_e$ or recombination in these microbes).

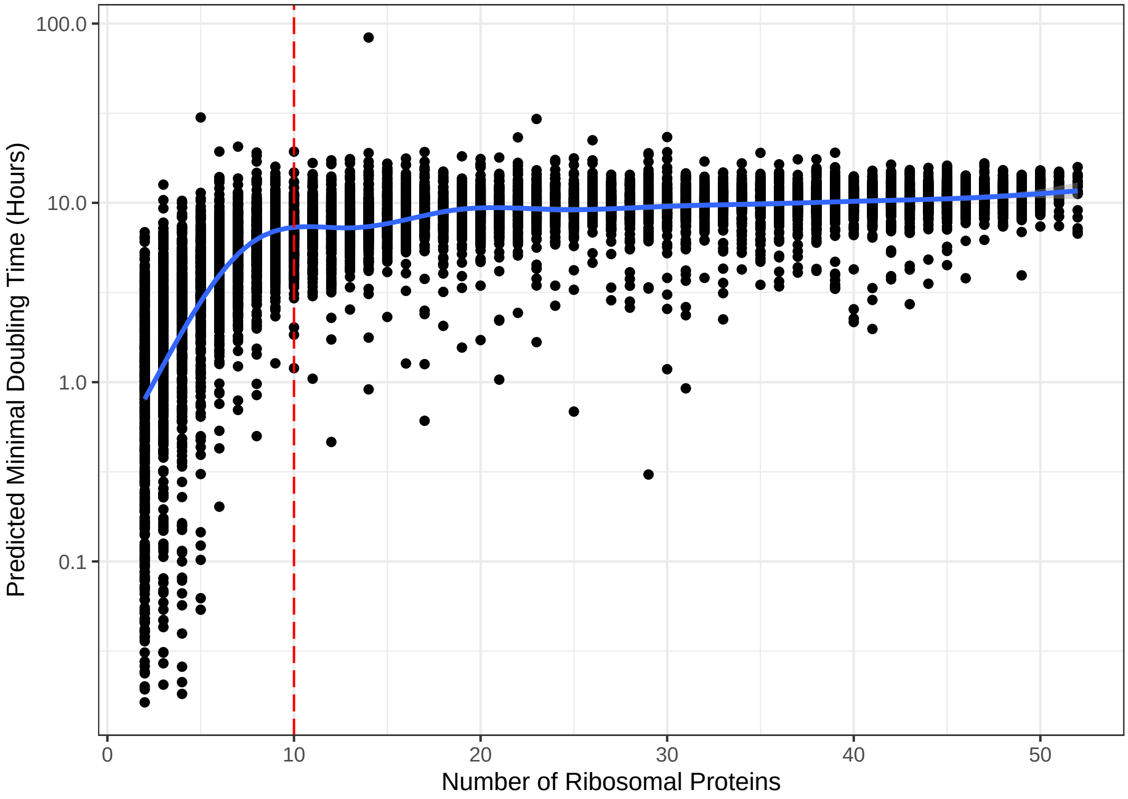


Figure S20: For SAGs with $< 10$ annotated ribosomal proteins growth rate predictions are biased. We do not include genomes below this threshold in any of our analyses.
